# Predicting antiviral resistance mutations in SARS-CoV-2 main protease with computational and experimental screening

**DOI:** 10.1101/2022.08.24.505060

**Authors:** Vishnu M. Sasi, Sven Ullrich, Jennifer Ton, Sarah E. Fry, Jason Johansen-Leete, Richard J. Payne, Christoph Nitsche, Colin J. Jackson

## Abstract

The main protease (M^pro^) of SARS-CoV-2 is essential for viral replication and has been the focus of many drug discovery efforts since the start of the COVID-19 pandemic. Nirmatrelvir (NTV) is an inhibitor of SARS-CoV-2 M^pro^ that is used in the combination drug Paxlovid for the treatment of mild to moderate COVID-19. However, with increased use of NTV across the globe, there is a possibility that future SARS-CoV-2 lineages will evolve resistance to NTV. Early prediction and monitoring of resistance mutations could allow for measures to slow the spread of resistance and for the development of new compounds with activity against resistant strains. In this work, we have used *in silico* mutational scanning and inhibitor docking of M^pro^ to identify potential resistance mutations. Subsequent *in vitro* experiments revealed five mutations (N142L, E166M, Q189E, Q189I, and Q192T) that reduce the potency of NTV and of a previously identified non-covalent cyclic peptide inhibitor of M^pro^. The E166M mutation reduced the half-maximal inhibitory concentration (IC_50_) of NTV 24-fold, and 118-fold for the non-covalent peptide inhibitor. Our findings inform the ongoing genomic surveillance of emerging SARS-CoV-2 lineages.

**Graphical Abstract:** 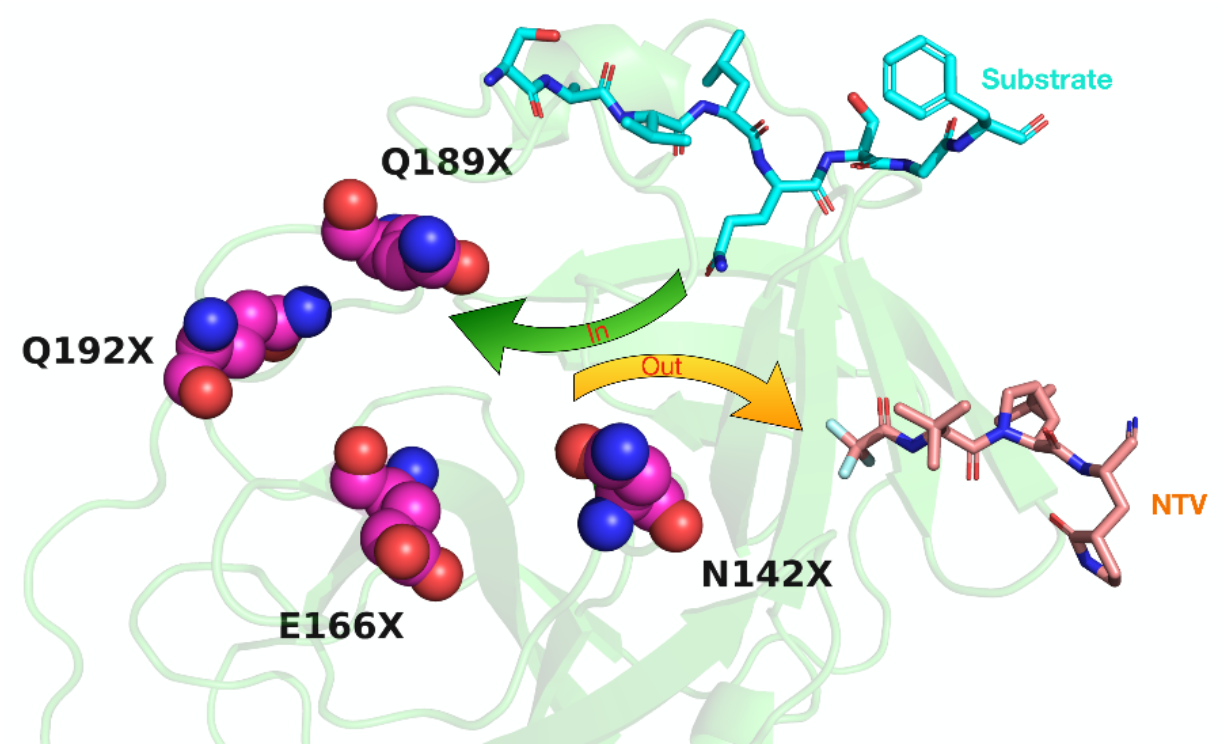

## Introduction

COVID-19, the disease resulting from infection by the coronavirus SARS-CoV-2, was declared a global pandemic by the World Health Organization (WHO) in March 2020.^1^ Since then, over 595 million infections have been recorded,^2^ with affected individuals experiencing symptoms ranging from fever and dry cough to life-threatening illness.^3^ Several lineages of SARS-CoV-2 have arisen, and five (Alpha, Beta, Gamma, Delta, Omicron) have been classified as variants of concern (VOC) by the WHO.^4^ Vaccines,^5^ monoclonal antibodies,^6^ and small molecule drugs,^7^ have all been rapidly developed and deployed clinically to aid in combating the COVID-19 pandemic.

The main protease (M^pro^, nsp5, 3CL^pro^; NCBI: YP_09742612) plays a critical role in the SARS-CoV-2 replication cycle by proteolytically liberating the majority of the non-structural proteins from the viral polyproteins (UniProt: P0DTC1, P0DTD1),^8^ some of which are involved in the replicase-transcription complex (RTC).^9^ M^pro^ assembles as a homodimer, whereby each of the protomers consists of three domains. Domain I (8–101) and domain II (102–184) form an antiparallel β-barrel structure and domain III (201–306) consists of a-helices arranged in an antiparallel globular cluster.^8, 10, 11^ The active site is located between domains I and II and includes the loop region (185–200) that connects the two domains.^8^ M^pro^ is a cysteine protease and employs a catalytic dyad (H41, C145) to catalyze hydrolysis of its substrates.^8^ The enzyme has a pronounced substrate specificity from P_4_ to P_1_’,^8^ using Schechter-Berger notation,^12^ whereby hydrolysis exclusively occurs after Q residues in P_1_.^13^ There is a preference for L in P_2_ and small residues such as A and S in P1’.^8, 13^ As no human protease possesses comparable peptide substrate specificity, M^pro^ is an ideal target for antiviral drug development.^8, 14^

Since the identification of SARS-CoV-2, a number of inhibitors of M^pro^ have been reported.^15^ While several non-covalent inhibitors are described,^16–18^ the field is dominated by covalent peptide-based inhibitors derived from the substrate sequence.^19^ As of August 2022, the covalent inhibitor nirmatrelvir (NTV)^20^ is the only approved M^pro^ inhibitor.^21^ NTV is a short, substrate-derived, peptidomimetic that binds to the catalytic cysteine through an electrophilic C-terminal nitrile warhead.^20^ *In vitro*, NTV possesses low nanomolar inhibition constants (*K*_i_) against the SARS-CoV-2 M^pro^ of the ancestral strain (in this study referred to as wild-type) and the Omicron variant.^22^

The anti-coronaviral drug Paxlovid contains both NTV and low-dose ritonavir (RTV), whereby the latter acts as pharmacokinetic booster of NTV.^21^ More than 12 million doses of Paxlovid have been manufactured and distributed to over 37 countries.^23^ With Paxlovid treatment now accessible to millions around the world, there is a high risk for current circulating SARS-CoV-2 variants to evolve for resistance to NTV. Drug resistance to protease inhibitors has previously been reported for several other infectious viral species,^24–26^ making it prudent to monitor the circulating SARS-CoV-2 variants for early identification of NTV resistance. Indeed, several recent preprints disclose experimentally found M^pro^ variants which reduce the potency of NTV.^27–33^ These studies encompassed a variety of methodologies, including data mining of SARS-CoV-2 sequences and passage experiments. Especially substitutions of substrate binding pocket residues such as S144, E166, L167, and Q192 resulted in loss of NTV inhibition (**Supporting Information Table 3**).

In this study we have used an *in silico* and experimental *in vitro* workflow for the identification of potential SARS-CoV-2 M^pro^ mutants that lose affinity for NTV, while maintaining substrate processing capability, i.e., mutations that could result in a viable resistant strain. Firstly, mutational scanning was performed on a selected number of residues in and around the enzyme’s active site to generate all possible single amino acid variant models. Covalent docking was then used to screen a substrate analogue and the inhibitor (NTV) against these mutants; mutations that significantly affected NTV binding and retained the ability to bind the substrate analogue were taken forward for experimental *in vitro* validation against NTV and an unrelated non-covalent cyclic peptide inhibitor. The resistant mutations identified in our study could be identified as mutations of interest before they evolve naturally, thereby informing the scientific and medical community ahead of time.

## Methods

### *In silico* protein and ligand preparation

The X-ray crystal structure of SARS-CoV-2 M^pro^ (PDB: 7VH8)^34^ was imported into Maestro (Schrödinger Release 2021-3: Maestro; Schrödinger, LLC: New York, NY, USA, 2021) and refined using the protein preparation wizard module with the addition of missing hydrogens and sidechains, converting selenomethionine to methionine, deleting water molecules, assigning bond orders, and generating heteroatom states at a pH of 7.4. The structure was further optimized using PROPKA at pH of 7.4 and minimized using OPLS4 forcefield,^35^ with the heavy atoms converged to RMSD of 0.3 Å. The structures of NTV and the substrate peptide were drawn in ChemDraw 20.0 (Perkin Elmer, USA) and optimized to generate 3D conformations using the LigPrep module in Maestro.

### M^pro^ mutant model generation

The M^pro^ X-ray crystal structure (PDB: 7VH8)^34^ was taken as a template to build models for Set 1-3 mutants using Residue Scanning Calculations (Schrödinger Release 2021-3: BioLuminate; Schrödinger, LLC: New York, NY, USA, 2021). All generated structures were refined with backbone minimisation using Prime at a cut-off distance of 5.0 Å around the mutated residue along with the prediction of stabilities parameters, namely: Δ stability (solvated, hydropathy, vdW, Hbond, Prime energy, etc.).

### Covalent docking

CovDock (Schrödinger Release 2021-3: Glide; Schrödinger, LLC: New York, NY, USA, 2021) was utilized for performing covalent docking of both the inhibitor and substrate against prepared mutants. For protomer A, the docking grid coordinates x = −19.2, y = 17.4, and z = −25.3 were utilized while for protomer B grid coordinates x = 19.2, y = 46.1, and z = −25.3 were found to be optimal for binding pose and docking score reproducibility. CovDock was performed using virtual screening mode and the number of output structures to be generated was set to 10. For the inhibitor docking, the reaction type was set to “Nucleophilic Addition to a Triple Bond” with the default SMARTS “[C]#[N]” selected, while for the substrate docking “Nucleophilic Addition to a Double Bond” was selected from the reaction type menu and the SMARTS “O=CC(N)CCC(=O)N” was manually added.

### Recombinant protein expression

Codon optimized genes encoding for the various SARS-CoV-2 M^pro^ variants were cloned into pET-29b(+) plasmid vectors between NdeI and XhoI restriction sites by Twist Bioscience (USA). The plasmids were transformed into *Escherichia coli* BL21 (DE3) and were plated onto a petri dish containing lysogeny broth (LB) media treated with 100 mg/L kanamycin (Sigma-Aldrich, USA). The seed culture was prepared by picking a single colony from the petri dish, which was inoculated with 10 mL LB media containing 100 mg/L kanamycin. The seed culture was incubated at 37 °C with shaking at 200 rpm for 18 h and was then diluted 1:100 into 1 L using the autoinduction media, terrific broth (TB) supplemented with lactose and 100 mg/L kanamycin. This was kept in a shaker (200 rpm) for 24 h at 20 °C, after which the cells were harvested (5000 *g* for 15 min at 4 °C) and stored at −20 °C.

### Protein purification

Cell pellets were resuspended in buffer A (50 mM Tris-HCl, 300 mM NaCl, 5 mM imidazole, pH 7.5) with the addition of 1 μL turbonuclease (Sigma-Aldrich, USA), followed by a two-cycle sonication (Omni International Mixer Homogenizer; Thermo Fisher, USA) for cell lysis. The suspension was centrifuged (13000 rpm for 30 min at 4 °C), and the resultant supernatant was first passed through a 0.45 μm filter followed by loading onto a His-Trap Ni^2+^ column (HisTrap FF; Cytiva, USA). The column was washed with buffer A and the protein was eluted using buffer B (50 mM Tris-HCl, 300 mM NaCl, 300 mM imidazole, pH 7.5). The protein fractions were analyzed using SDS-PAGE and the pure fractions were combined and concentrated, with a molecular weight cut-off at 10 kDa, using the exchange buffer, buffer C (20 mM HEPES-KOH, 150 mM NaCl, 1 mM DTT, 1 mM EDTA, pH 7.0). The protein concentrations were determined using a UV/vis-spectrophotometer (NanoDrop One^C^, Thermo Fisher, USA) by measuring the absorbance at 280 nm using the extinction coefficients (M^-1^cm^-1^ at 280 nm) as calculated with ExPASy ProtParam. ^36^

### Enzyme inhibition assays

The SARS-CoV-2 M^pro^ FRET assay was carried out as previously described.^18, 37^ Triplicate enzymatic reactions (100 μL) were prepared in black polypropylene 96-well plates with U-bottom (Greiner Bio-One, Austria) and monitored by the microplate reader Infinite 200 PRO M Plex (Tecan, Switzerland). 20 mM Tris-HCl pH 7.3, 100 mM NaCl, 1 mM EDTA, 1 mM DTT was used as the aqueous buffer. Nirmatrelvir (MedChemExpress, USA; Batch HY-138687-116180) was incubated with recombinant SARS-CoV-2 M^pro^ (final enzyme concentration: 25 nM) for 10 min at 37 °C, before the fluorogenic substrate (*DABCYL*)-KTSAVLQ↓SGFRKME(*EDAN5*)-NH_2_ (Mimotopes, Australia) was added to a concentration of 25 μM to initiate the reaction (whereby *DABCYL* and *EDANS* act as the FRET pair and ↓ denotes the proteolytic cleavage site). The enzymatic activity was monitored at 37 °C for 5 min at 490 nm using an excitation wavelength of 340 nm. For IC_50_ determination from dose-response curves, the initial velocities of the control reactions were defined as 100 % activity to calculate inhibition percentages accordingly. The data were analyzed and visualized with Prism 9.4 (GraphPad Software, USA), using the sigmoidal four-parameter logistic curve model with plateaus at 0 and 100 %.

## Results

### Computational screening of potential resistance mutations

A total of 711 mutant models of SARS-CoV-2 M^pro^ were generated *in silico* for the study in three sets (**Supporting Information Table 1**). Set 1 consisted of residues lining the substrate/inhibitor binding site. To select these residues, we first analyzed the substrate and inhibitor bound SARS-CoV-2 M^pro^ structures (PDB: 7TE0 and 7N89, respectively).^38–40^ Set 1 includes the sub-pockets S_4_–S_1_ and S_1_’–S_3_’ at the active site, which contribute to both substrate and inhibitor non-covalent interactions with the residues P_4_-P_1_ and P_1_’-P_3_’ respectively, in addition to other residues within van der Waals, salt bridge or hydrogen bonding distance to substrate or inhibitor. The specific interactions between enzyme and its various peptide recognition sites have recently been mapped, and are consistent with this initial set (T25, T26, L27, H41, S46, M49, Y54, F140, L141, N142, G143, S144, H163, H164, M165, E166, L167, P168, F185, D187, Q189, T190, A191, and Q192) (**Figure 1, Supporting Information Figure 1**).^38^ Because NTV is significantly smaller than the substrate, several of these residues are beyond non-covalent interaction distance to it (e.g., Thr25–Leu27), but were retained in Set 1 regardless, given the possibility that structural disruption could still affect inhibitor binding. These were mutated to all possible single point mutations *in silico,* yielding 456 mutant structures. For the second set, residues situated proximal to the binding site, namely M130, R131, G146, S147, C160, T169, G170, V171, H172, P184, V186, A193 and A194 were selected and mutated to all possible amino acids to generate an additional 247 mutant structures (**Figure 1, Supporting Information Figure 1**). Residues M130, R131, G146 and S147 form part of the active site loop 129-148, while C160, T169, G170 and H172 belong to the anti-parallel β-sheet 156-175 lining the binding site. Residues P184, V186, A193 and A194 were chosen because they form a part of the loop 176-200, which connects domain II to domain III and is located close to the active site of M^pro^. The third set of mutations consisted of eight currently circulating SARS-CoV-2 M^pro^ mutants G15S, T21I, Q83K, L89F, K90R, P132H, L205V, and V212F (**Supporting Information Figure 1**),^37, 41^ which were selected to assess whether mutations at these positions could affect substrate or NTV binding.

**Figure 1.**
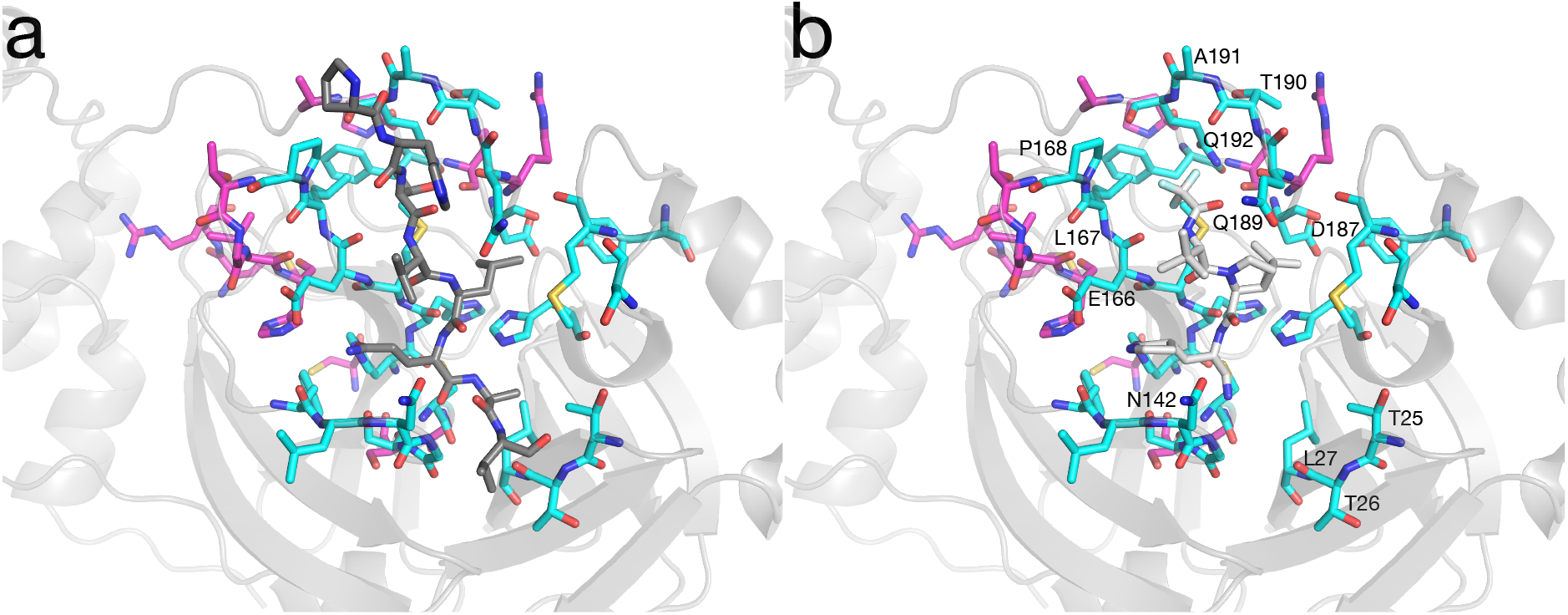
Locations of *in silico* mutated sites in M^pro^. *In silico* mutagenesis was used to generate 711 mutants in three sets: (i) Set 1 residues lining the substrate (a; SAVLQSGF) or NTV (b) (grey sticks) binding site (cyan sticks; T25, T26, L27, H41, S46, M49, Y54, F140, L141, N142, G143, S144, H163, H164, M165, E166, L167, P168, F185, D187, Q189, T190, A191 and Q192); (ii) Set 2 residues in the second shell of the binding site and in functionally important loops (magenta sticks; M130, R131, G146, S147, C160, T169, G170, V171, H172, P184, V186, A193 and A194).

Having generated an *in silico* library of 711 SARS-CoV-2 M^pro^ mutants, we then sought to assess whether the mutations would affect binding of either the native substrate or the inhibitor NTV (or both). Using CovDock,^42^ we performed an initial screen against the substrate; this is important if a mutation is to be viable as a resistance mutation, i.e., it must not affect the native activity to such a degree that it has a substantial fitness cost. This is especially true in this case because the SARS-CoV-2 M^pro^ enzyme is essential and there is little redundancy; if the mutation abolishes SARS-CoV-2 M^pro^ function, the virus will not be viable. SARS-CoV-2 M^pro^ can hydrolyze a variety of related polypeptide sequences and is somewhat promiscuous.^38^ Given that this was an initial step, and that it is more likely the docking screen will underestimate the number of deleterious mutations (i.e., be more likely to yield false positives),^43, 44^ and thus will be relatively permissive and unlikely to erroneously screen out mutations that would retain activity versus the native substrate(s), we used only one sequence, SAVLQSGF. The purpose of this step was to reduce the library size by eliminating mutations that would clearly abolish native activity (and result in a fitness loss). The results of this initial screen reveal that a significant number of these mutations result in reduced binding of a representative native substrate (**Figure 2**). Interestingly, the number of deleterious mutations from Set 2 was comparable to that from Set 1, most likely because of indirect structural disruptions rather than direct effects on the substrate. From this first screen, the number of mutations of interest was narrowed down from 711 to 513. As expected, the circulating mutants all retained wild-type like substrate binding (**Supporting Information Figure 1**).

**Figure 2.**
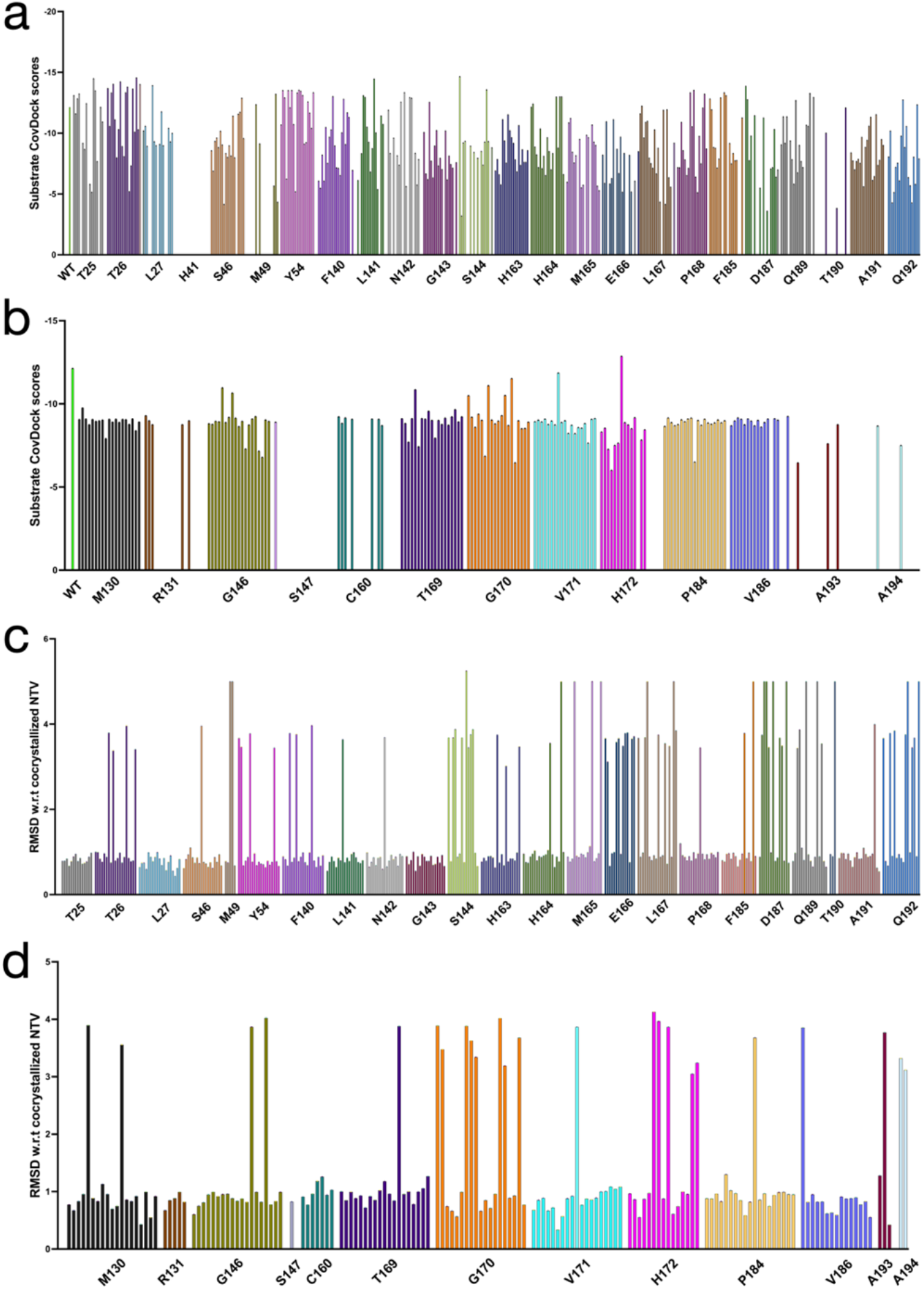
Computational screening of M^pro^ mutant structures. (a) Docking scores for the substrate (SAVLQSGF) against the modelled M^pro^ mutants from Set 1. (b) Substrate docking scores plotted for the Set 2 mutant library. (c) RMSD plot of the docked inhibitor (NTV), from the selected Set 1 mutants, with respect to the co-crystallized NTV pose in WT M^pro^. (d) RMSD of the docked inhibitor (NTV), from the selected Set 2 mutants, with respect to the cocrystallized NTV pose in WT M^pro^. All the mutants have been sorted into groups of 19 for each residue position and a docking score of 0 is used to represent mutants producing no binding scores against the substrate.

Mutants that potentially retain native activity from the substrate screen were then screened with the inhibitor, NTV, again using CovDock. The docking scores themselves were not particularly informative and likely include many false positives (**Supporting Information Table 1**). Thus, instead of relying on the docking score, we calculated the inhibitor root mean square deviation (RMSD) relative to the crystal structure of bound NTV to better estimate the disruption to NTV binding (**Figure 2**). This was more informative, providing a greater difference between mutants where binding was not affected and those where it potentially was. This indicated that, while many of the mutants are predicted to retain sensitivity to NTV, a subset of these mutations (111) is predicted to significantly disrupt NTV binding. As expected, none of the currently circulating variants (Set 3) resulted in significant disruption to NTV binding (**Supporting Information Figure 1)**.

### Medium throughput expression and native activity screening

Of the 111 potential resistance mutations identified through the computational screen, we decided to experimentally verify 32, to test for the effects of the mutations on expression, native activity, and inhibition by NTV. These were selected by choosing all candidate resistance variants identified from the NTV CovDock screen from 9 positions: N142(L), E166(A/C/L/M/P/T), L167(A/E/F/M/W/Y), P168(M), D187(F/H/N/S/T/W), Q189(D/E/I/P/T), T190(P), A191(V), and Q192(C/H/P/T/Y). These 9 positions were chosen on the basis of a combination of proximity to NTV (**Figure 1**), high RMSD (e.g., D187, T190, etc.; **Figure 2**), and to ensure the mutations were distributed across all key binding sub-sites of the inhibitor-protein complex (**Figure 1)**. This reduced dataset allowed us to test the hypothesis that resistance mutations can arise at the substrate binding site and allow for the identification of a limited (non-comprehensive) number of potential resistance mutations. Accordingly, genes for the N142L, E166A/C/L/M/P/T, L167A/E/F/M/W/Y, P168M, D187F/H/N/S/T/W, Q189D/E/I/P/T, T190P, A191V, and Q192C/H/P/T/Y variants were synthesized and cloned into an expression vector (pET-29b(+)) for heterologous expression and purification from *Escherichia coli*.

The effect of a mutation on organismal fitness depends on the expression of the function of the gene, e.g., through enzyme activity. However, this first requires that the gene that is expressed is properly translated and the resulting protein folded into its stable and active native tertiary structure. Thus, the first step in this medium throughput experimental screen tested for soluble protein expression. From this, it was observed that seven of these variants did not express in soluble form: E166P, L167E, L167M, L167W, D187H, D187N, and Q192Y. This is somewhat consistent with the predicted effects of these mutations on protein stability, as all of these were predicted to be destabilizing (**Supporting Information Table 2**). Given that the protein expression was performed in a recombinant organism, we cannot be certain that these mutations will also result in impaired protein expression in the host cell during viral replication. However, from the combination of the predicted destabilizing effects and the lack of expression, which generally correlates with low stability of the folded protein or of a folding intermediate,^45, 46^ we can conclude they have a lower likelihood of folding to the mature, active form.

The remaining expressed and soluble mutants were then screened against a substrate analogue (*DABCYL*)-KTSAVLQSGFRKME(*EDANS*)-NH_2_) in which the peptide recognition sequence is sandwiched between 5-((2-aminoethyl)amino)naphthalene-1-sulfonic acid (*EDANS*) and 4-((4-(dimethylamino)phenyl)azo)benzoic acid (*DABCYL*), for which cleavage results in a loss of Förster resonance energy transfer (FRET) that can be detected spectroscopically (**Figure 3**). This revealed that several of the mutants (E166A/C/L/T, L167A/F/Y, D187F/S/T/W/D, Q189P/T, T190P, A191V, Q192H) had no detectable activity with this substrate analogue at the measured concentration. While this initially appears to be a larger than expected number of inactive variants, it is consistent with the generally high rate of false positive “hits” from docking, i.e., docking experiments tend to overestimate the binding of ligands. It could well be that the mutations sample conformations that block binding in solution that were not captured in the computational docking. Interestingly, two of these mutations have previously been reported to have WT-like activity *in vitro* (Q189P)^29^ or in a vesicular stomatitis virus-based cellular assay (L167F),^30^ although the functional assays used in those studies differed from that used here. We also cannot discount the possibility that the enzymes lost activity during purification or storage. Of the 8 active variants, four (E166M, P168M, Q192C, Q192T) exhibited a modest reduction in activity of no more than ~5-fold, whilst three variants (N142L, Q189E, Q189I) exhibited greater than wild-type activity. The Q192P variant exhibited a more pronounced reduction in activity (~10 fold) as compared to the wild-type. Thus, from the medium throughput screen, we identified seven assumed neutral and one near-neutral (Q192P) mutations to SARS-CoV-2 M^pro^ that could be sufficiently neutral in terms of expression, stability, and activity that could accumulate in SARS-CoV-2 without major impairment of fitness.

**Figure 3.**
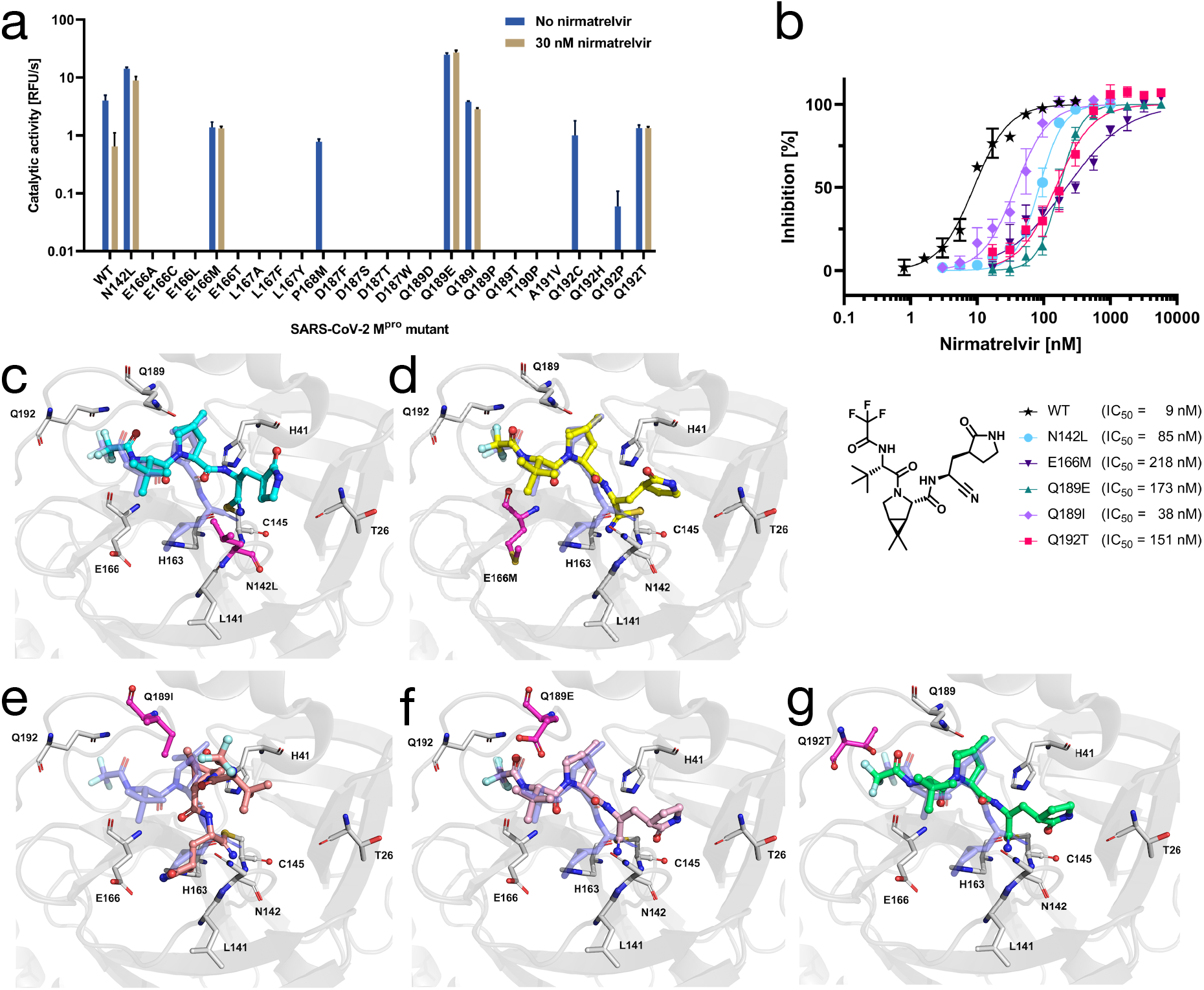
(a) Experimental analysis of the activity of 25 SARS-CoV-2 M^pro^ mutants with fluorogenic substrate alone and in the presence of NTV (30 nM). Several of the mutants were not active with this substrate analog, suggesting they are likely to have significant effects on fitness. Two mutants remained sensitive to NTV inhibition (P168M, Q192C and Q192P). Five mutants (N142L, E166M, Q189E, Q189I, Q192T) displayed reduced inhibition by NTV. (b) Dose-response curves of NTV against SARS-CoV-2 M^pro^ mutants. (c–g) Structural effects of the mutations (N142L, E166M, Q189E, Q189I, Q192T in that order) showing disruption to the binding of NTV compared with binding of NTV to WT M^pro^ (grey).

### Screening of neutral variants against NTV

The inhibition of this final set of eight neutral/near-neutral mutants by NTV was then tested using single-point assays (30 nM; approximately 3-fold higher than the wild-type IC_50_ measured in this study) (**Figure 3, Supporting Information Table 4**). This revealed that, at this concentration, where wild-type was inhibited to ~15% original activity, the P168M, Q192P and Q192C were all at least as well inhibited as the wild-type. In these variants, the activity with NTV present fell below the limit of detection (consistent with their uninhibited activity starting from a lower baseline than wildtype). However, NTV appeared to be substantially less effective against five M^pro^ variants (N142L, E166M, Q189E, Q189I, Q192T). To quantify the effect of these mutations on the inhibitory activity of NTV, we determined the half-maximal inhibitory concentrations (IC_50_) of NTV against these mutants (**Figure 3**). All five IC_50_ values were higher than those of NTV against the WT protease in our assay conditions (IC_50_ = 9 nM), with the largest difference between WT and E166M (24-fold, IC_50_ = 218 nM). Mutations Q189E (IC_50_ = 173 nM) and Q192T (IC_50_ = 151 nM) were the next most impactful mutations, reducing the IC_50_ 19-fold and 17-fold, respectively. Substitutions N142L (IC_50_ = 85 nM) and Q189I (IC_50_ = 38 nM) caused 9-fold and 4-fold IC_50_ reductions. All five mutations are predicted to significantly disrupt NTV binding (**Figure 3c–g**). Thus, at least five mutations to residues near to the binding site of NTV have the potential to retain stable folding, native activity (and thus likely have minimal fitness cost) and to also reduce the effectiveness of NTV.

### Testing the effects of resistance mutations on a non-covalent macrocyclic peptide inhibitor

To evaluate the effects of these mutations on an alternative inhibitor, we tested an unrelated macrocyclic peptide inhibitor (“Peptide 1”), which was discovered from a Random non-standard Peptides Integrated Discovery (RaPID)^47^ mRNA display selection (**Figure 4**).^18^ In contrast to most peptide-based inhibitors of M^pro^,^15^ including NTV, Peptide 1 displays nanomolar affinity without a covalent warhead. The inhibition of wild-type SARS-CoV-2 M^pro^ with Peptide 1 (IC_50_ = 60 nM) in this study was in line with the literature value.^18^ We then tested the inhibition of the potential resistance mutants N142L, E166M, Q189E, Q189I, and Q192T by Peptide 1 to assess the effects of these mutations on a non-covalent inhibitor that also has affinity in the nanomolar range (**Figure 4, Supporting Information Table 4**). These results reveal interesting and heterogeneous trends in terms of the inhibition relative to NTV. E166M was substantially more effective at reducing inhibition by peptide 1, increasing the IC_50_ 118-fold (*vs.* 24-fold for NTV); Q189I also had a greater effect at reducing inhibition (19-fold *vs.* 4-fold for NTV). In contrast, N142L, Q189E and Q192T all exhibited only minor changes in inhibition (1.5–5.5 fold) in comparison to their effects on NTV inhibition (9–19-fold). Structural analysis is consistent with these results (**Figure 4**): E166 interacts with the critical P1 glutamine (Q3 in Peptide 1), Q189 likewise forms a hydrogen bond with the main chain NH of the L2 and would be more affected by a Q-I mutation than Q-E, while N142 hydrogen is in a pocket where hydrophobic interactions with the thioether resulting from the N-L mutation could compensate for lost hydrogen bonds, and Q192 is comparatively remote from the inhibitor. These results suggest that potential resistance mutations are likely to have relatively specific effects with different compounds, raising the possibility that combination therapy could reduce the chances of resistance developing.^46^

**Figure 4.**
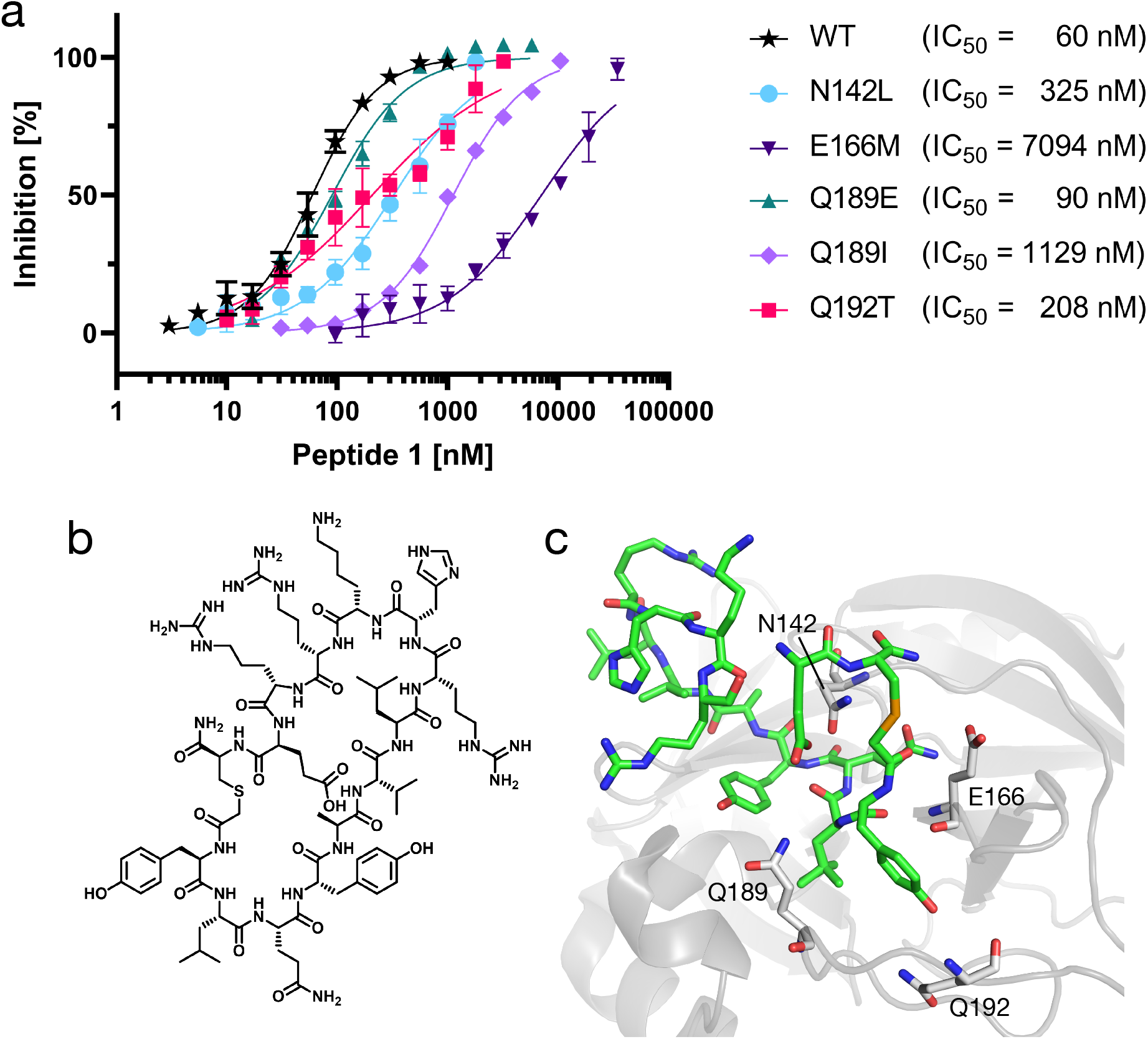
Effects of potential resistance mutations on the inhibition of SARS-CoV-2 M^pro^ by a non-covalent peptide inhibitor. (a) Structure of Peptide 1. (b) Dose-response curves of peptide 1 against SARS-CoV-2 M^pro^ mutants. (c) Structure of the Peptide 1: M^pro^ complex (modelled from PDB: 7RNW)^18^ showing the positions of the potential resistance mutations (N142, E166, Q189, Q192) relative to Peptide 1. Note that the thioether in Peptide 1 is a selenoether in the co-crystal structure. P2 leucine is shown in multiple side chain conformations.

### Screening of sequence databases for potential resistance mutations

To understand whether any of the five M^pro^ mutations identified here (N142L, E166M, Q189E, Q189I, Q192T) were already in circulation, we consulted the EpiCov database by GISAID.^48^ As of 8^th^ August 2022, professional contributors had uploaded over 12 million SARS-CoV-2 sequences on the platform. We detected 177 complete SARS-CoV-2 sequences with the previously discussed M^pro^ amino acid substitutions (**Figure 5**, **Supporting Information Table 5**; N142L: 20, E166M: 1, Q189E: 11, Q189I: 6, Q192T: 139). Of these 177 sequences, 72 were collected after 22^nd^ December 2021, the emergency approval date of Paxlovid by the FDA (N142L: 1, Q189E: 10, Q189I: 6, Q192T: 55). To put these numbers into perspective, we also calculated the number of sequences containing the Omicron signature M^pro^ mutation P132H,^22, 37^ of which more than 5 million sequences are accessible. Given the low number of deposited sequences with our five identified mutations, we conclude that SARS-CoV-2 viruses containing these M^pro^ mutations do not circulate in relevant numbers at the time of writing. Finally, we compared the number of mutations (with >75% prevalence) in SARS-CoV-2 M^pro^ with the number of mutations in the spike protein (NCBI: YP_009724390). This shows that considerably more mutations have been fixed in the evolution of spike (31) *vs.* M^pro^ (1), suggesting that while spike has been under selective pressure to adapt and evolve, M^pro^ is evolving comparatively slowly, without significant accumulation of neutral mutations, which is discussed in more detail later.

**Figure 5.**
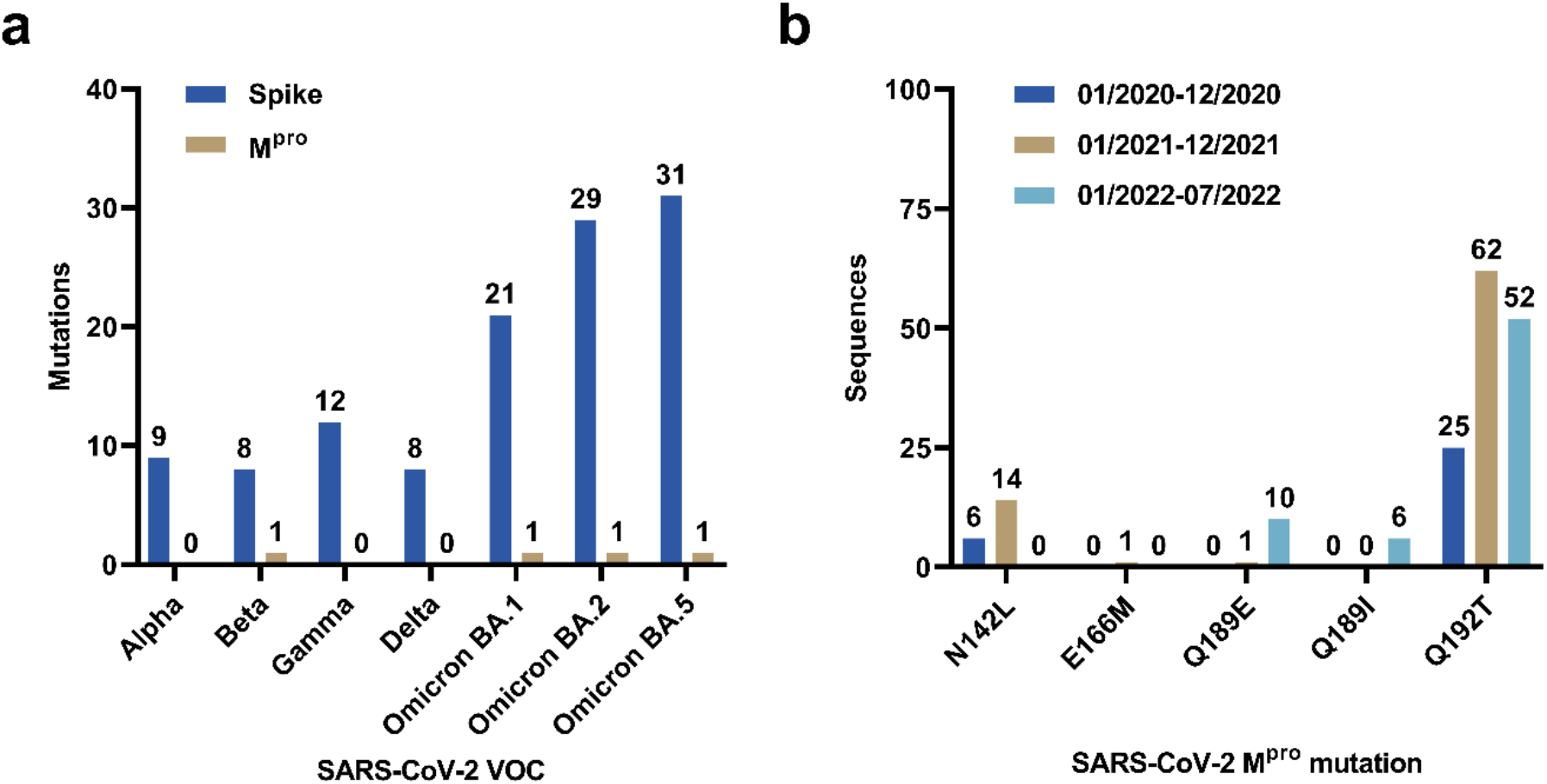
(a) Spike and M^pro^ mutations (> 75 % prevalence) in different SARS-CoV-2 variants of concern lineages (according to WHO). The data were accessed via the Outbreak.info lineage comparison report.^52^ (b) SARS-CoV-2 sequences accessed via the GISAID EpiCoV database containing M^pro^ mutations N142L, E166M, Q189E, Q189I or Q192T by date of sample collection.

A final consideration regarding the risk of evolution of resistance is whether these mutations are accessible via single nucleotide changes. The M^pro^ amino acids N142, E166, Q189 and Q192 are encoded in the viral RNA as AAU (N), GAA (E), and CAA (Q), respectively (WIV04 reference sequence).^49^ The E166M mutation (GAA → AUG) requires three mutations from the WIV04 reference sequence, while the N142L substitution requires two point mutations (AUU → UUA/UUG). The Q189E mutation only requires one (CAA → GAA) or two (CAA → GAG) point mutations. Both the Q189I mutations require two (CAA → AUA) or three (CAA → AUU/AUC) point mutations, as does the Q192T mutation (CAA → ACA or CAA → ACU/ACC/ACG). It has been discussed that such mutations in the SARS-CoV-2 genome may have implications for the secondary structure of the viral RNA,^50, 51^ which raises the question of whether additional selective pressures might be involved in the evolution of resistance against NTV.

## Discussion

During our study, multiple preprints have appeared, discussing certain SARS-CoV-2 M^pro^ mutations and their effect on NTV.^27–33^ The methodologies used in these studies to discover the mutations with resistance potential are diverse, which is reflected in their results. As is observed in our study, previously reported mutational hotspots include the active site of M^pro^, with special emphasis on the residues S144, E166, L167, and Q192. While N142L did not decrease the potency of NTV in one study,^29^ it was found to be ~9-fold less susceptible to NTV than the WT enzyme here. The same study reported that Q192T decreases the IC_50_ of NTV ~7-fold, which correlates well with our results (~17-fold decrease).^29^ Other mutations of E166 reduced the potency of NTV in other studies,^27–29^ consistent with our observation that E166M (not studied prior to this work) potentially has dramatic effects on NTV binding. The importance of E166 is likely related to a major hydrogen bond between M^pro^ and NTV in P1.^20^ Mutant Q189E was reported to increase the IC_50_ value of NTV by up to ~2-fold,^29^ while we found a ~19-fold increase. While Q189 is a prominent residue to trigger NTV resistance,^29^ our study is the first to report that the mutation Q189I affects NTV inhibition. In summary, our study independently confirms findings reported by other groups and adds new data to the pool of mutations that reduce the inhibition of NTV (e.g., E166M, Q189I). Interestingly, we found stronger effects for certain mutants than previously reported (e.g., Q192T, Q189E), highlighting the importance of verification of results across multiple laboratories and methodologies.

In a study such as this, it is important to identify the limitations, so that the results are not interpreted out of context. The first obvious limitation of our study is that neither viral fitness nor actual drug resistance of the discovered mutants can be demonstrated in *in vivo* models, as this would require potentially dangerous gain-of-function experiments with SARS-CoV-2 that our laboratories lack the capacity to perform. Secondly, our computational screening and experimental assay were only based on one substrate recognition sequence. While the core recognition site that M^pro^ cleaves is well conserved,^8^ there is variation across the terminal regions of multiple recognition sites that M^pro^ cleaves.^38^ Thus, while it is likely that differences in the edges of the recognition sites will not be as critical to substrate recognition, it is possible that mutations we expect to be “neutral” with respect to the native function could lose activity with other untested substrate recognition sites. Unfortunately, the combinatorial complexity of the number of M^pro^ cleavage sites prevented us from excluding this possibility. Finally, this study (like others) is not comprehensive: we show that the computational pre-screen is an effective and economical method to reduce the library size for screening, yet we cannot exclude the probability that this initial high throughput screen could have missed additional mutations that could cause resistance. Indeed, we did not identify mutations that were discovered using alternative approaches (and *vice versa*). Thus, this study should be viewed as providing support for the hypothesis that resistance mutations, which can reduce NTV binding without significantly affecting the native activity, could occur, and to contribute to the other studies on the topic thereby increasing the number of mutations that should be monitored.

Our study is also unique in testing the potential evolution of resistance to both covalent and non-covalent inhibitors. The major benefits of covalent enzyme inhibition are reduced dependence on substrate competition and increased efficiency.^53^ It is therefore no surprise that the majority of high-affinity peptide-based M^pro^ inhibitors like NTV contain covalent modifiers, while only very few compounds, e.g. Peptide 1 (**Figure 4**), can achieve similar affinity without a covalent warhead.^18^ Both inhibitors, NTV (e.g., PDB: 7VH8, 7RFW, 7TLL)^20, 22, 34^ and peptide 1 (PDB: 7RNW),^18^ have been co-crystallized with M^pro^. One interesting comparison is the effect of the E166M mutation, which has reduced interactions (relative to E166) with either glutamine or a glutamine mimetic in P1. The binding of Peptide 1 is reduced 5-times more than the binding of NTV due to this mutation (a similar reduction of affinity is apparent for the Q189E mutation). A plausible explanation for this is that the covalent attachment of NTV to M^pro^ provides some robustness to the development of resistance. However, we also see that other mutants have less of an effect, which can be rationalized by their reduced interactions with Peptide 1, relative to NTV. These results suggest that it could be possible to adapt inhibitors such as NTV to some degree to avoid or mitigate the loss of binding caused by some mutations, or to develop combination therapies in which a single mutation cannot reduce binding of two different inhibitors.

The implication of these results on the evolutionary processes underlying the emergence of resistance can also be discussed. The lack of genetic diversity in the M^pro^ gene is initially surprising, given the mutation rate of SARS-CoV-2 and the comparatively rapid accumulation of mutations in the spike protein. Likewise, there is relatively little neutral genetic drift apparent in available sequences, although several of these mutations have been detected previously at very low frequency (**Figure 5**). While these results appear to be encouraging, the explanation for this observation is that SAR-CoV-2 has been evolving in a series of selective “sweeps”,^54^ in which strains with enhanced immune evasion and virulence rapidly become dominant and, in the process, displace other strains and reduce genetic diversity overall (e.g. neutral genetic variation in other genes such as that which encode M^pro^). We should not assume that this will continue forever; the longer a strain is dominant the more we will observe genetic variation in other genes such as that encoding M^pro^, raising the chance that a variant with selective advantage (e.g., resistance to NTV) will emerge and spread. We identify some mutations here that are easily accessible through single nucleotide changes, which are likely to emerge rapidly, and others that require two or three mutations. Given the mutation rate of SARS-CoV-2, and the likelihood that intermediate mutants (e.g., E166Q on the way to E166M)^29^ could also confer resistance to a lower level, it should not be assumed that the requirement for multiple mutations will provide a long-term protection once resistance to Paxlovid is a strong selective pressure.

To summarize, we report five SARS-CoV-2 M^pro^ variants (N142L, E166M, Q189E, Q189I, Q192T) that are associated with reduced NTV efficacy *in vitro*. The most-impactful mutation in terms of their effect on NTV potency were E166M (IC_50_ = 218 nM, 24-fold change) and Q189E (IC_50_ = 173 nM, 19-fold change). Both E166 and Q189 were previously found to be involved in hydrogen bond formation with NTV.^20^ The substitutions had ~5-times greater effect on the binding of a non-covalent peptide inhibitor. At the time of writing, the five identified mutations N142L, E166M, Q189E, Q189I, and Q192T are not observed at significant frequency in circulating SARS-CoV-2 strains, and several of these will require multiple nucleotide changes to be accessed. However, this highlights the importance of genome surveillance (especially of “breakthrough” infections), so that newly circulating lineages with resistance to NTV can be identified as soon as possible. Despite the availability of two small molecule drugs against COVID-19, potential drug resistance highlights that it continues to be important to identify and progress more antivirals, to eventually increase the number of available SARS-CoV-2 therapeutics.

## Acknowledgments

C.N. thanks the Australian Research Council (ARC) for a Discovery Early Career Research Award (DE190100015) and Discovery Project funding (DP200100348). C.J.J., C.N. and R.J.P. acknowledge support by the ARC Centre of Excellence for Innovations in Peptide & Protein Science (CE200100012) and the ARC Centre of Excellence in Synthetic Biology (CE200100029). This study was supported by a RAMR (MAWA) grant awarded to S.U. and C.N. We thank Dr. Elwy H. Abdelkader (Australian National University) for assistance with protein mass spectrometry. We thank A/Prof. Rob Lanfear (Australian National University) for helpful discussions on SARS-CoV-2 sequence data. We gratefully acknowledge all SARS-CoV-2 sequence data contributors, i.e., the authors and their originating laboratories responsible for obtaining the specimens, and their submitting laboratories for generating the genetic sequence and metadata and sharing via GISAID, on which parts of this research are based.

## References

(1) Mahase, E. (2020) COVID-19: WHO declares pandemic because of “alarming levels” of spread, severity, and inaction, BMJ 368, m1036.

(2) Dong, E., Du, H., and Gardner, L. (2020) An interactive web-based dashboard to track COVID-19 in real time, Lancet Infect. Dis. 20, 533–534.

(3) Osuchowski, M. F., Winkler, M. S., Skirecki, T., Cajander, S., Shankar-Hari, M., Lachmann, G., Monneret, G., Venet, F., Bauer, M., Brunkhorst, F. M., Weis, S., Garcia-Salido, A., Kox, M., Cavaillon, J.-M., Uhle, F., Weigand, M. A., Flohé, S. B., Wiersinga, W. J., Almansa, R., de la Fuente, A., Martin-Loeches, I., Meisel, C., Spinetti, T., Schefold, J. C., Cilloniz, C., Torres, A., Giamarellos-Bourboulis, E. J., Ferrer, R., Girardis, M., Cossarizza, A., Netea, M. G., van der Poll, T., Bermejo-Martín, J. F., and Rubio, I. (2021) The COVID-19 puzzle: deciphering pathophysiology and phenotypes of a new disease entity, Lancet Respir. Med. 9, 622–642.

(4) Telenti, A., Hodcroft, E. B., and Robertson, D. L. (2022) The evolution and biology of SARS-CoV-2 variants, Cold Spring Harb. Perspect. Med. 12, a041390.

(5) Simões, R. S. d. Q., and Rodríguez-Lázaro, D. (2022) Classical and next-generation vaccine platforms to SARS-CoV-2: biotechnological strategies and genomic variants, Int. J. Environ. Res. Public Health 19, 2392.

(6) Evans, M. J., Kumar, S., Chandele, A., and Sharma, A. (2021) Current status of therapeutic monoclonal antibodies against SARS-CoV-2, PLOS Pathog. 17, e1009885.

(7) Saravolatz, L. D., Depcinski, S., and Sharma, M. (2022) Molnupiravir and nirmatrelvir-ritonavir: oral COVID antiviral drugs, Clin. Infect. Dis., ciac180.

(8) Ullrich, S., and Nitsche, C. (2020) The SARS-CoV-2 main protease as drug target, Bioorg. Med. Chem. Lett. 30, 127377.

(9) Hartenian, E., Nandakumar, D., Lari, A., Ly, M., Tucker, J. M., and Glaunsinger, B. A. (2020) The molecular virology of coronaviruses, J. Biol. Chem. 295, 12910–12934.

(10) Roe, M. K., Junod, N. A., Young, A. R., Beachboard, D. C., and Stobart, C. C. (2021) Targeting novel structural and functional features of coronavirus protease nsp5 (3CLpro, Mpro) in the age of COVID-19, J. Gen. Virol. 102, 001558.

(11) Jin, Z., Du, X., Xu, Y., Deng, Y., Liu, M., Zhao, Y., Zhang, B., Li, X., Zhang, L., Peng, C., Duan, Y., Yu, J., Wang, L., Yang, K., Liu, F., Jiang, R., Yang, X., You, T., Liu, X., Yang, X., Bai, F., Liu, H., Liu, X., Guddat, L. W., Xu, W., Xiao, G., Qin, C., Shi, Z., Jiang, H., Rao, Z., and Yang, H. (2020) Structure of Mpro from SARS-CoV-2 and discovery of its inhibitors, Nature 582, 289–293.

(12) Schechter, I., and Berger, A. (1967) On the size of the active site in proteases. I. Papain, Biochem. Biophys. Res. Commun. 27, 157–162.

(13) Rut, W., Groborz, K., Zhang, L., Sun, X., Zmudzinski, M., Pawlik, B., Wang, X., Jochmans, D., Neyts, J., Młynarski, W., Hilgenfeld, R., and Drag, M. (2020) SARS-CoV-2 Mpro inhibitors and activity-based probes for patient-sample imaging, Nat. Chem. Biol. 17, 222–228.

(14) Hilgenfeld, R. (2014) From SARS to MERS: crystallographic studies on coronaviral proteases enable antiviral drug design, FEBS J. 281, 4085–4096.

(15) Lv, Z., Cano, K. E., Jia, L., Drag, M., Huang, T. T., and Olsen, S. K. (2022) Targeting SARS-CoV-2 proteases for COVID-19 antiviral development, Front. Chem. 9, 819165.

(16) Kitamura, N., Sacco, M. D., Ma, C., Hu, Y., Townsend, J. A., Meng, X., Zhang, F., Zhang, X., Ba, M., Szeto, T., Kukuljac, A., Marty, M. T., Schultz, D., Cherry, S., Xiang, Y., Chen, Y., and Wang, J. (2021) Expedited approach toward the rational design of noncovalent SARS-CoV-2 main protease inhibitors, J. Med. Chem. 65, 2848–2865.

(17) Unoh, Y., Uehara, S., Nakahara, K., Nobori, H., Yamatsu, Y., Yamamoto, S., Maruyama, Y., Taoda, Y., Kasamatsu, K., Suto, T., Kouki, K., Nakahashi, A., Kawashima, S., Sanaki, T., Toba, S., Uemura, K., Mizutare, T., Ando, S., Sasaki, M., Orba, Y., Sawa, H., Sato, A., Sato, T., Kato, T., and Tachibana, Y. (2022) Discovery of S-217622, a noncovalent oral SARS-CoV-2 3CL protease inhibitor clinical candidate for treating COVID-19, J. Med. Chem. 65, 6499–6512.

(18) Johansen-Leete, J., Ullrich, S., Fry, S. E., Frkic, R., Bedding, M. J., Aggarwal, A., Ashhurst, A. S., Ekanayake, K. B., Mahawaththa, M. C., Sasi, V. M., Luedtke, S., Ford, D. J., O’Donoghue, A. J., Passioura, T., Larance, M., Otting, G., Turville, S., Jackson, C. J., Nitsche, C., and Payne, R. J. (2022) Antiviral cyclic peptides targeting the main protease of SARS-CoV-2, Chem. Sci. 13, 3826–3836.

(19) Ullrich, S., Sasi, V. M., Mahawaththa, M. C., Ekanayake, K. B., Morewood, R., George, J., Shuttleworth, L., Zhang, X., Whitefield, C., Otting, G., Jackson, C., and Nitsche, C. (2021) Challenges of short substrate analogues as SARS-CoV-2 main protease inhibitors, Bioorg. Med. Chem. Lett. 50, 128333.

(20) Owen, D. R., Allerton, C. M. N., Anderson, A. S., Aschenbrenner, L., Avery, M., Berritt, S., Boras, B., Cardin, R. D., Carlo, A., Coffman, K. J., Dantonio, A., Di, L., Eng, H., Ferre, R., Gajiwala, K. S., Gibson, S. A., Greasley, S. E., Hurst, B. L., Kadar, E. P., Kalgutkar, A. S., Lee, J. C., Lee, J., Liu, W., Mason, S. W., Noell, S., Novak, J. J., Obach, R. S., Ogilvie, K., Patel, N. C., Pettersson, M., Rai, D. K., Reese, M. R., Sammons, M. F., Sathish, J. G., Singh, R. S. P., Steppan, C. M., Stewart, A. E., Tuttle, J. B., Updyke, L., Verhoest, P. R., Wei, L., Yang, Q., and Zhu, Y. (2021) An oral SARS-CoV-2 Mpro inhibitor clinical candidate for the treatment of COVID-19, Science 374, 1586–1593.

(21) Lamb, Y. N. (2022) Nirmatrelvir plus ritonavir: first approval, Drugs 82, 585–591.

(22) Greasley, S. E., Noell, S., Plotnikova, O., Ferre, R., Liu, W., Bolanos, B., Fennell, K., Nicki, J., Craig, T., Zhu, Y., Stewart, A. E., and Steppan, C. M. (2022) Structural basis for the in vitro efficacy of nirmatrelvir against SARS-CoV-2 variants, J. Biol. Chem. 298, 101972.

(23) Pfizer. (2022). Pfizer to invest $120 million to produce COVID-19 oral treatment in the US, https://www.pfizer.com/news/press-release/press-release-detail/pfizer-invest-120-million-produce-covid-19-oral-treatment.

(24) Mason, S., Devincenzo, J. P., Toovey, S., Wu, J. Z., and Whitley, R. J. (2018) Comparison of antiviral resistance across acute and chronic viral infections, Antivir. Res. 158, 1031–12.

(25) Wensing, A. M. J., van Maarseveen, N. M., and Nijhuis, M. (2010) Fifteen years of HIV protease inhibitors: raising the barrier to resistance, Antivir. Res. 85, 59–74.

(26) Martinez, M. A., and Franco, S. (2020) Therapy implications of hepatitis C virus genetic diversity, Viruses 13, 41.

(27) Zhou, Y., Gammeltoft, K. A., Ryberg, L. A., Pham, L. V., Fahnøe, U., Binderup, A., Hernandez, C. R. D., Offersgaard, A., Fernandez-Antunez, C., Peters, G. H. J., Ramirez, S., Bukh, J., and Gottwein, J. M. (2022) Nirmatrelvir resistant SARS-CoV-2 variants with high fitness in vitro, bioRxiv, 2022.2006.2006.494921.

(28) Jochmans, D., Liu, C., Donckers, K., Stoycheva, A., Boland, S., Stevens, S. K., De Vita, C., Vanmechelen, B., Maes, P., Trüeb, B., Ebert, N., Thiel, V., De Jonghe, S., Vangeel, L., Bardiot, D., Jekle, A., Blatt, L. M., Beigelman, L., Symons, J. A., Raboisson, P., Chaltin, P., Marchand, A., Neyts, J., Deval, J., and Vandyck, K. (2022) The substitutions L50F, E166A and L167F in SARS-CoV-2 3CLpro are selected by a protease inhibitor *in vitro* and confer resistance to nirmatrelvir, bioRxiv, 2022.2006.2007.495116.

(29) Hu, Y., Lewandowski, E. M., Tan, H., Morgan, R. T., Zhang, X., Jacobs, L. M. C., Butler, S. G., Mongora, M. V., Choy, J., Chen, Y., and Wang, J. (2022) Naturally occurring mutations of SARS-CoV-2 main protease confer drug resistance to nirmatrelvir, bioRxiv, 2022.2006.2028.497978.

(30) Heilmann, E., Costacurta, F., Volland, A., and von Laer, D. (2022) SARS-CoV-2 3CLpro mutations confer resistance to Paxlovid (nirmatrelvir/ritonavir) in a VSV-based, non-gain-of-function system, bioRxiv, 2022.2007.2002.495455.

(31) de Oliveira, V. M., Ibrahim, M. F., Sun, X., Hilgenfeld, R., and Shen, J. (2022) H172Y mutation perturbs the S1 pocket and nirmatrelvir binding of SARS-CoV-2 main protease through a nonnative hydrogen bond, bioRxiv, 2022.2007.2031.502215.

(32) Iketani, S., Mohri, H., Culbertson, B., Hong, S. J., Duan, Y., Luck, M. I., Annavajhala, M. K., Guo, Y., Sheng, Z., Uhlemann, A.-C., Goff, S. P., Sabo, Y., Yang, H., Chavez, A., and Ho, D. D. (2022) Multiple pathways for SARS-CoV-2 resistance to nirmatrelvir, bioRxiv, 2022.2008.2007.499047.

(33) Moghadasi, S. A., Heilmann, E., Moraes, S. N., Kearns, F. L., von Laer, D., Amaro, R. E., and Harris, R. S. (2022) Transmissible SARS-CoV-2 variants with resistance to clinical protease inhibitors, bioRxiv, 2022.2008.2007.503099.

(34) Zhao, Y., Fang, C., Zhang, Q., Zhang, R., Zhao, X., Duan, Y., Wang, H., Zhu, Y., Feng, L., Zhao, J., Shao, M., Yang, X., Zhang, L., Peng, C., Yang, K., Ma, D., Rao, Z., and Yang, H. (2021) Crystal structure of SARS-CoV-2 main protease in complex with protease inhibitor PF-07321332, Protein Cell 13, 689–693.

(35) Lu, C., Wu, C., Ghoreishi, D., Chen, W., Wang, L., Damm, W., Ross, G. A., Dahlgren, M. K., Russell, E., Von Bargen, C. D., Abel, R., Friesner, R. A., and Harder, E. D. (2021) OPLS4: improving force field accuracy on challenging regimes of chemical space, J. Chem. Theory Comp. 17, 4291–4300.

(36) Gasteiger, E., Gattiker, A., Hoogland, C., Ivanyi, I., Appel, R. D., and Bairoch, A. (2003) ExPASy: the proteomics server for in-depth protein knowledge and analysis, Nucleic Acids Res. 31, 3784–3788.

(37) Ullrich, S., Ekanayake, K. B., Otting, G., and Nitsche, C. (2022) Main protease mutants of SARS-CoV-2 variants remain susceptible to nirmatrelvir, Bioorg. Med. Chem. Lett. 62, 128629.

(38) Shaqra, A. M., Zvornicanin, S. N., Huang, Q. Y. J., Lockbaum, G. J., Knapp, M., Tandeske, L., Bakan, D. T., Flynn, J., Bolon, D. N. A., Moquin, S., Dovala, D., Kurt Yilmaz, N., and Schiffer, C. A. (2022) Defining the substrate envelope of SARS-CoV-2 main protease to predict and avoid drug resistance, Nat. Commun. 13, 3556.

(39) Yang, K. S., Leeuwon, S. Z., Xu, S., and Liu, W. R. (2022) Evolutionary and structural insights about potential SARS-CoV-2 evasion of nirmatrelvir, J. Med. Chem. 65, 8686–8698.

(40) Kneller, D. W., Zhang, Q., Coates, L., Louis, J. M., and Kovalevsky, A. (2021) Michaelis-like complex of SARS-CoV-2 main protease visualized by room-temperature X-ray crystallography, IUCrJ 8, 973–979.

(41) Sacco, M. D., Hu, Y., Gongora, M. V., Meilleur, F., Kemp, M. T., Zhang, X., Wang, J., and Chen, Y. (2022) The P132H mutation in the main protease of Omicron SARS-CoV-2 decreases thermal stability without compromising catalysis or small-molecule drug inhibition, Cell Res. 32, 498–500.

(42) Zhu, K., Borrelli, K. W., Greenwood, J. R., Day, T., Abel, R., Farid, R. S., and Harder, E. (2014) Docking covalent inhibitors: a parameter free approach to pose prediction and scoring, J. Chem. Inf. Model. 54, 1932–1940.

(43) Perola, E. (2006) Minimizing false positives in kinase virtual screens, Proteins 64, 422–435.

(44) Cerón-Carrasco, J. P. (2022) When virtual screening yields inactive drugs: dealing with false theoretical friends, ChemMedChem, e202200278.

(45) Jackson, C. J., Coppin, C. W., Carr, P. D., Aleksandrov, A., Wilding, M., Sugrue, E., Ubels, J., Paks, M., Newman, J., Peat, T. S., Russell, R. J., Field, M., Weik, M., Oakeshott, J. G., Scott, C., and Kivisaar, M. (2014) 300-fold increase in production of the Zn2+-dependent dechlorinase TrzN in soluble form via apoenzyme stabilization, Appl. Environ. Microbiol. 80, 4003–4011.

(46) Sugrue, E., Scott, C., and Jackson, C. J. (2017) Constrained evolution of a bispecific enzyme: lessons for biocatalyst design, Org. Biomol. Chem. 15, 937–946.

(47) Passioura, T., and Suga, H. (2017) A RaPID way to discover nonstandard macrocyclic peptide modulators of drug targets, Chem. Commun. 53, 1931–1940.

(48) Khare, S., Gurry, C., Freitas, L., B Schultz, M., Bach, G., Diallo, A., Akite, N., Ho, J., Tc Lee, R., Yeo, W., Core Curation Team, G., and Maurer-Stroh, S. (2021) GISAID’s role in pandemic response, China CDC Weekly 3, 1049–1051.

(49) Zhou, P., Yang, X.-L., Wang, X.-G., Hu, B., Zhang, L., Zhang, W., Si, H.-R., Zhu, Y., Li, B., Huang, C.-L., Chen, H.-D., Chen, J., Luo, Y., Guo, H., Jiang, R.-D., Liu, M.-Q., Chen, Y., Shen, X.-R., Wang, X., Zheng, X.-S., Zhao, K., Chen, Q.-J., Deng, F., Liu, L.-L., Yan, B., Zhan, F.-X., Wang, Y.-Y., Xiao, G.-F., and Shi, Z.-L. (2020) A pneumonia outbreak associated with a new coronavirus of probable bat origin, Nature 579, 270–273.

(50) Lan, T. C. T., Allan, M. F., Malsick, L. E., Woo, J. Z., Zhu, C., Zhang, F., Khandwala, S., Nyeo, S. S. Y., Sun, Y., Guo, J. U., Bathe, M., Näär, A., Griffiths, A., and Rouskin, S. (2022) Secondary structural ensembles of the SARS-CoV-2 RNA genome in infected cells, Nat. Commun. 13, 1128.

(51) Hosseini Rad Sm, A., and McLellan, A. D. (2020) Implications of SARS-CoV-2 mutations for genomic RNA structure and host microRNA targeting, Int. J. Mol. Sci. 21, 4807.

(52) Gangavarapu, K., Abdel Latif, A., Mullen, J. L., Alkuzweny, M., Hufbauer, E., Tsueng, G., Haag, E., Zeller, M., Aceves, C. M., Zaiets, K., Cano, M., Zhou, J., Qian, Z., Sattler, R., Matteson, N. L., Levy, J. I., Suchard, M. A., Wu, C., Su, A. I., Andersen, K. G., and Hughes, L. D. (2022) Outbreak.info genomic reports: scalable and dynamic surveillance of SARS-CoV-2 variants and mutations, medRxiv, 2022.2001.2027.22269965.

(53) Bauer, R. A. (2015) Covalent inhibitors in drug discovery: from accidental discoveries to avoided liabilities and designed therapies, Drug Discov. Today 20, 1061–1073.

(54) Kang, L., He, G., Sharp, A. K., Wang, X., Brown, A. M., Michalak, P., and Weger-Lucarelli, J. (2021) A selective sweep in the Spike gene has driven SARS-CoV-2 human adaptation, Cell 184, 4392–4400.e4394.

