## Supplementary Material for "Predicting antiviral resistance mutations in SARS-CoV-2 main protease with computational and experimental screening"

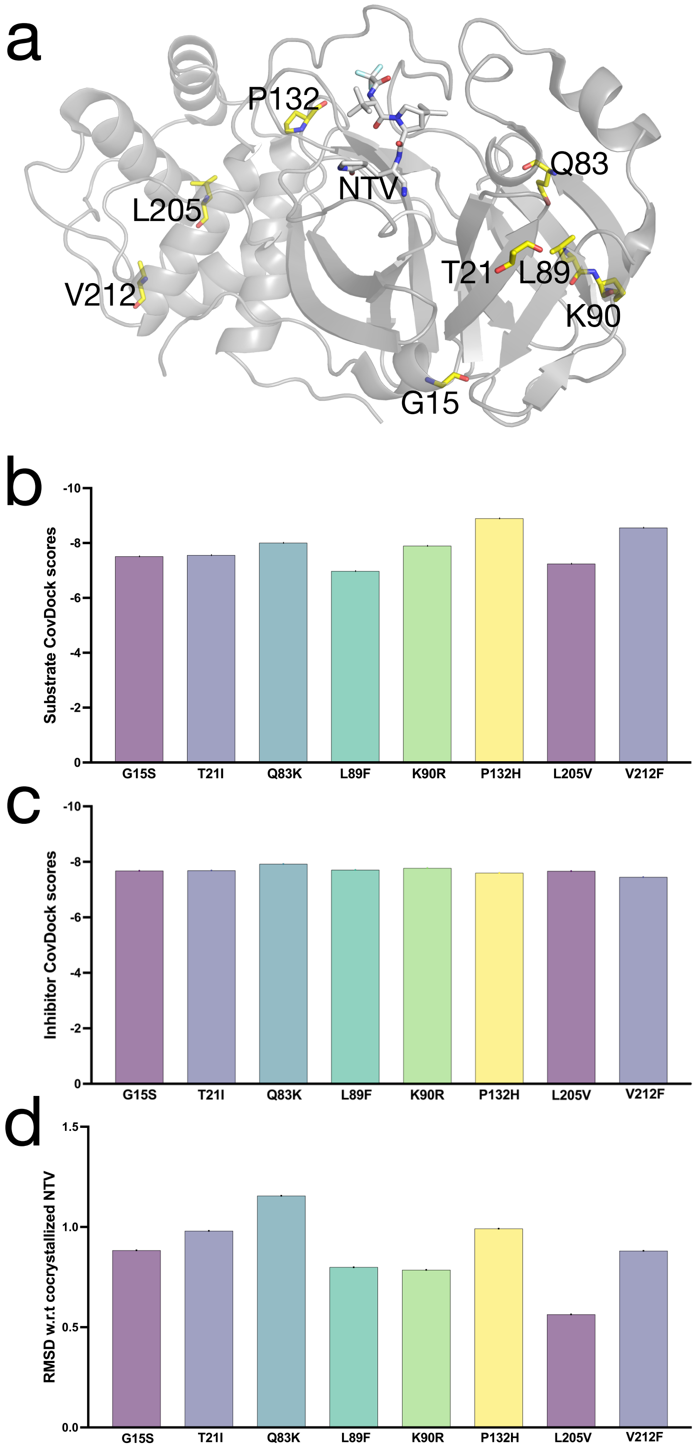


**Supporting Information Figure 1.** Currently circulating $M^{pro}$ sequence polymorphisms. (a) Location of the mutations in the $M^{pro}$ monomer (PDB: 7VH8).[1] (b–d) Results from computational docking experiments showing little effect on binding of substrate (b) or NTV (c–d).

**Supporting Information Table 2.** Predicted effects of the 111 potential resistance mutations that passed the screens for maintenance of substrate binding and disruption of NTV binding on thermostability.

| Mutant | Δ Stability (solvated) [kJ/mol] |
|---|---|
| T26A | 11.36 |
| T26D | 22.46 |
| T26H | 12.23 |
| T26K | 17.55 |
| T26R | -9.44 |
| T26Y | 13.57 |
| S46E | 0.04 |
| S46K | 18.48 |
| M49V | 18.15 |
| M49W | 22.01 |
| Y54A | 58.29 |
| Y54C | 55.66 |
| Y54G | 73.64 |
| Y54T | 49.26 |
| M130C | 50.14 |
| F140E | 51.12 |
| F140I | 55.16 |
| F140R | 3.29 |
| L141K | 7.56 |
| N142L | -9.1 |
| S144A | 0.78 |
| S144D | 82.64 |
| S144E | 20.36 |
| S144K | 42.69 |
| S144M | 62.58 |
| S144P | 108.42 |
| S144Q | 7.34 |
| S144R | -31.03 |
| S144T | 16.06 |
| G146F | 386.78 |
| G146T | 123.83 |
| H163C | 25.5 |
| H163D | 64.04 |
| H163F | 12.41 |
| H163I | 11.65 |
| H163K | 19.98 |

| | |
|---|---|
| H163M | -12.02 |
| H163P | 153.48 |
| H163T | 36.91 |
| H163W | 55.14 |
| H163Y | -10.7 |
| H164N | 20.89 |
| H164Q | -11.01 |
| H164W | 191.8 |
| M165F | 42.9 |
| M165W | 47.29 |
| M165Y | 26.17 |
| E166A | 18.64 |
| E166C | 14.7 |
| E166L | 5.79 |
| E166M | -13.48 |
| E166P | 30.16 |
| E166N | 9.01 |
| E166T | 19.88 |
| E166W | 11.13 |
| E166Y | 7.09 |
| L167A | 45.43 |
| L167E | 36.16 |
| L167F | 221.94 |
| L167M | 7.29 |
| L167Q | 27.41 |
| L167T | 33.27 |
| L167W | 287.68 |
| P168M | -0.89 |
| T169F | 6.99 |
| G170A | 2.93 |
| G170E | 25.55 |
| G170H | 58.85 |
| G170I | 5.55 |
| G170P | 40.05 |
| G170R | -8.75 |
| G170S | 4.08 |
| G170Y | 6.08 |
| V171I | -8.74 |
| H172E | 47.75 |
| H172I | 24.35 |
| H172Q | -29.07 |

| | |
|---|---|
| H172W | 65.21 |
| H172Y | 50.59 |
| P184L | -0.32 |
| F185Q | 16.92 |
| F185W | 33.89 |
| V186A | 10.69 |
| D187C | 6.45 |
| D187E | 35.59 |
| D187F | 98.02 |
| D187H | 50.7 |
| D187N | 9.76 |
| D187S | 8.85 |
| D187T | 10.78 |
| D187W | 121.41 |
| Q189D | 16.98 |
| Q189E | 2.93 |
| Q189I | -6.89 |
| Q189P | 52.14 |
| Q189T | 6.31 |
| T190P | 122.97 |
| A191V | -0.17 |
| Q192C | 30.7 |
| Q192F | 23.69 |
| Q192H | 32.73 |
| Q192P | 109.69 |
| Q192R | 5.83 |
| Q192T | 24.64 |
| Q192V | 16.32 |
| Q192Y | 164.8 |
| A192C | -2.41 |
| A193D | 5.86 |
| A193N | 2.65 |
| A194I | 2.04 |
| A194R | -2.94 |

**Supporting Information Table 3.** SARS-CoV-2 $M^{pro}$ mutations corroborated to affect NTV potency and the respective bioRxiv preprint authors.

| SARS-CoV-2 mutations | Preprint study |
|---|---|
| T21I<br>T21I+T304I<br>L50F<br>L50F+E166V<br>L50F+A173V<br>E166V<br>T304 | Zhou et al.[2] |
| L50F+E166A<br>L50F+E166A+L167F<br>E166A<br>L167F | Jochmans et al.[3] |
| S144A/F/G/M/Y<br>M165T<br>E166Q<br>H172F/Q<br>Q192S/T/V | Hu et al.[4] |
| Y54C<br>L167F<br>L167F+F305L<br>Q192R<br>Q192R+F305L | Heilmann et al.[5] |
| H172Y | de Oliveira et al.[6] |
| T45I<br>D48Y<br>S144A<br>ΔP168<br>A173T<br>A173V<br>ΔP168+A173V | Iketani et al.[7] |
| T21I<br>L50F<br>S144A<br>E166V<br>A173V<br>P252L<br>T304I<br>T21I+S144A<br>T21I+E166V<br>T21I+A173V<br>T21I+T304I<br>L50F+E166V<br>T21I+A173V+T304I | Moghadasi et al.[8] |

**Supporting Information Table 4.** $IC_{50}$ values of NTV and Peptide 1 against SARS-CoV-2 $M^{pro}$ variants with 95% confidence intervals shown in brackets.

| SARS-CoV-2 variant | NTV $IC_{50}$ [nM] | Peptide 1 $IC_{50}$ [nM] |
|---|---|---|
| WT | 9 (8–10) | 60 (55–65) |
| N142L | 85 (80–91) | 325 (281–376) |
| E166M | 218 (182–262) | 7094 (6108–8245) |
| Q189E | 173 (163–183) | 90 (81–101) |
| Q189I | 38 (33–43) | 1129 (1071–1190) |
| Q192T | 151 (128–179) | 208 (170–256) |

**Supporting Information Table 5.** List of SARS-CoV-2 sequences with $M^{pro}$ mutations N142, E166M, Q189E, Q189I or Q192T compared to the ancestral strain reference genome (hCoV-19/Wuhan/WIV04/2019; EPI_ISL_402124).[9] The sequences were extracted from the EpiCoV database of the GISAID[10] (complete sequences, low coverage excluded) on 8th August 2022. The collection date format is YYYY-MM-DD.

| $M^{pro}$ mutation | Virus sequence name | GISAID accession ID | Collection date |
|---|---|---|---|
| N142L | hCoV-19/India/GJ-GBRC20/2020 | EPI_ISL_437449 | 2020-04-26 |
| N142L | hCoV-19/India/GJ-GBRC39/2020 | EPI_ISL_444470 | 2020-04-29 |
| N142L | hCoV-19/India/GJ-GBRC78a/2020 | EPI_ISL_447536 | 2020-05-05 |
| N142L | hCoV-19/India/GJ-GBRC79a/2020 | EPI_ISL_447538 | 2020-05-05 |
| N142L | hCoV-19/India/GJ-GBRC79b/2020 | EPI_ISL_447539 | 2020-05-05 |
| N142L | hCoV-19/Slovenia/08-035564-MB/2021 | EPI_ISL_1056582 | 2021-02-04 |
| N142L | hCoV-19/India/GJ-ICMR-23/2020 | EPI_ISL_1164933 | 2020-05 |
| N142L | hCoV-19/Slovenia/17-020093-CE/2021 | EPI_ISL_1240615 | 2021-02-24 |
| N142L | hCoV-19/USA/AZ-ASU2815/2021 | EPI_ISL_1364772 | 2021-02-11 |
| N142L | hCoV-19/USA/AZ-ASU2941/2021 | EPI_ISL_1365537 | 2021-02-16 |
| N142L | hCoV-19/USA/AZ-ASU3694/2021 | EPI_ISL_1592329 | 2021-03-14 |
| N142L | hCoV-19/USA/ID-IBL-725613/2021 | EPI_ISL_3217201 | 2021-04-26 |
| N142L | hCoV-19/USA/ID-IBL-746167/2021 | EPI_ISL_3588737 | 2021-08-04 |
| N142L | hCoV-19/USA/ID-IBL-746524/2021 | EPI_ISL_3588800 | 2021-08-03 |
| N142L | hCoV-19/Germany/BW-RKI-I-267558/2021 | EPI_ISL_4933747 | 2021-09-02 |
| N142L | hCoV-19/Germany/BW-RKI-I-267649/2021 | EPI_ISL_4933831 | 2021-09-02 |
| N142L | hCoV-19/England/MILK-28BEA14/2021 | EPI_ISL_5994871 | 2021-10-29 |
| N142L | hCoV-19/USA/ID-IBL-767393/2021 | EPI_ISL_6340469 | 2021-10-19 |
| N142L | hCoV-19/USA/ID-IBL-770144/2021 | EPI_ISL_7314067 | 2021-11-09 |
| N142L | hCoV-19/USA/ID-IBL-780842/2021 | EPI_ISL_8681664 | 2021-12-30 |
| E166M | hCoV-19/USA/TX-HMH-MCoV-59197/2021 | EPI_ISL_5346845 | 2021-08-27 |
| Q189E | hCoV-19/USA/ID-IBL-748688/2021 | EPI_ISL_4211476 | 2021-08-12 |
| Q189E | hCoV-19/Italy/ABR-CAST-T2508/2022 | EPI_ISL_9135521 | 2022-01-17 |
| Q189E | hCoV-19/USA/LA-OD-1600491110/2022 | EPI_ISL_9570718 | 2022-01-24 |
| Q189E | hCoV-19/England/PHEC-5X07CZA8/2022 | EPI_ISL_11314407 | 2022 |
| Q189E | hCoV-19/England/PHEC-5X07DZ96/2022 | EPI_ISL_11314416 | 2022 |
| Q189E | hCoV-19/England/PHEC-5X084Z45/2022 | EPI_ISL_11376286 | 2022 |
| Q189E | hCoV-19/England/PHEC-6M0B5Z4F/2022 | EPI_ISL_12203945 | 2022 |
| Q189E | hCoV-19/Germany/HH-RKI-I-769476/2022 | EPI_ISL_12523646 | 2022-04-21 |
| Q189E | hCoV-19/USA/NY-PBRI-C22MY016/2022 | EPI_ISL_12605119 | 2022-05-02 |

| | | | |
|---|---|---|---|
| Q189E | hCoV-19/Germany/HH-RKI-I-858309/2022 | EPI_ISL_13266562 | 2022-05-19 |
| Q189E | hCoV-19/Pakistan/PPHRL-PACP-133/2022 | EPI_ISL_13728999 | 2022-06-28 |
| Q189I | hCoV-19/Spain/AS-242246223/2022 | EPI_ISL_10688175 | 2022-02-22 |
| Q189I | hCoV-19/England/PHEC-5W06AZ45/2022 | EPI_ISL_10925707 | 2022 |
| Q189I | hCoV-19/England/PHEC-5X077ZE3/2022 | EPI_ISL_11223119 | 2022 |
| Q189I | hCoV-19/England/PHEC-5X083ZA2/2022 | EPI_ISL_11376277 | 2022 |
| Q189I | hCoV-19/India/TN-SPHL-ICMR-INSACOG-0809/2022 | EPI_ISL_11887834 | 2022-01-17 |
| Q189I | hCoV-19/USA/LA-OD-5239691/2022 | EPI_ISL_13374815 | 2022-06-07 |
| Q192T | hCoV-19/Australia/VIC5550/2020 | EPI_ISL_10560677 | 2020-07-21 |
| Q192T | hCoV-19/Australia/VIC830/2020 | EPI_ISL_10559959 | 2020-04-04 |
| Q192T | hCoV-19/England/204661600/2020 | EPI_ISL_693485 | 2020-11-11 |
| Q192T | hCoV-19/England/PHEC-2FEBC6/2021 | EPI_ISL_2739679 | 2021-02-23 |
| Q192T | hCoV-19/England/PHEC-3Z07BZAB/2021 | EPI_ISL_8517275 | 2021 |
| Q192T | hCoV-19/England/PHEC-4G0BBZ2F/2021 | EPI_ISL_8738691 | 2021-12-14 |
| Q192T | hCoV-19/England/PHEC-4G0BBZ3E/2021 | EPI_ISL_8738695 | 2021 |
| Q192T | hCoV-19/England/PHEC-4W079Z38/2022 | EPI_ISL_9687048 | 2022 |
| Q192T | hCoV-19/England/PHEC-5P086Z30/2022 | EPI_ISL_9835937 | 2022 |
| Q192T | hCoV-19/England/PHEC-5U045Z28/2022 | EPI_ISL_10325873 | 2022 |
| Q192T | hCoV-19/England/PHEC-5V082Z96/2022 | EPI_ISL_10671808 | 2022 |
| Q192T | hCoV-19/England/PHEC-5V085Z42/2022 | EPI_ISL_10671863 | 2022 |
| Q192T | hCoV-19/England/PHEC-5V089ZEC/2022 | EPI_ISL_10757797 | 2022 |
| Q192T | hCoV-19/England/PHEC-5W031ZDA/2022 | EPI_ISL_10882516 | 2022 |
| Q192T | hCoV-19/England/PHEC-5W04EZ4B/2022 | EPI_ISL_10882642 | 2022 |
| Q192T | hCoV-19/England/PHEC-5X073ZB2/2022 | EPI_ISL_11223076 | 2022 |
| Q192T | hCoV-19/England/PHEC-5X07CZA8/2022 | EPI_ISL_11314407 | 2022 |
| Q192T | hCoV-19/England/PHEC-5X07DZ96/2022 | EPI_ISL_11314416 | 2022 |
| Q192T | hCoV-19/England/PHEC-5X084Z45/2022 | EPI_ISL_11376286 | 2022 |
| Q192T | hCoV-19/England/PHEC-5X08AZF8/2022 | EPI_ISL_11469618 | 2022 |
| Q192T | hCoV-19/England/PHEC-6M0B5Z03/2022 | EPI_ISL_12203944 | 2022 |
| Q192T | hCoV-19/England/PHEC-6M0B5Z4F/2022 | EPI_ISL_12203945 | 2022 |
| Q192T | hCoV-19/France/ARA-CHUGA-1065851800/2021 | EPI_ISL_2900249 | 2021-06-27 |
| Q192T | hCoV-19/Gibraltar/205121254/2020 | EPI_ISL_769872 | 2020 |
| Q192T | hCoV-19/India/MH-ICMR-NIV-INSACOG-GSEQ-9005/2022 | EPI_ISL_9969210 | 2022-01-31 |
| Q192T | hCoV-19/Pakistan/PPHRL-PACP-122/2022 | EPI_ISL_13728988 | 2022-01-26 |
| Q192T | hCoV-19/Pakistan/PPHRL-PACP-132/2022 | EPI_ISL_13728998 | 2022-06-28 |

| | | | |
|---|---|---|---|
| Q192T | hCoV-19/Spain/CT-HUGTiPM062EP6C4/2021 | EPI_ISL_7900442 | 2021-12-10 |
| Q192T | hCoV-19/Spain/MD-HGUGM-603284/2021 | EPI_ISL_2782337 | 2021-06-21 |
| Q192T | hCoV-19/USA/AZ-ASPHL-2377/2020 | EPI_ISL_2695941 | 2020-07-30 |
| Q192T | hCoV-19/USA/AZ-ASPHL-5247/2021 | EPI_ISL_4553009 | 2021-09-13 |
| Q192T | hCoV-19/USA/AZ-ASPHL-639/2020 | EPI_ISL_2227568 | 2020-05-08 |
| Q192T | hCoV-19/USA/CA-CZB-26882/2021 | EPI_ISL_2658594 | 2021-02-02 |
| Q192T | hCoV-19/USA/ID-IBL-763972/2021 | EPI_ISL_7950396 | 2021-10-12 |
| Q192T | hCoV-19/USA/LA-OD-3283875228/2022 | EPI_ISL_13104214 | 2022-05-23 |
| Q192T | hCoV-19/USA/LA-OD-5092500676/2022 | EPI_ISL_13104224 | 2022-05-25 |
| Q192T | hCoV-19/USA/LA-OD-9457230721/2021 | EPI_ISL_7856453 | 2021-12-13 |
| Q192T | hCoV-19/USA/MT-TRACE-GALL-100720-779/2020 | EPI_ISL_6938239 | 2020-10-07 |
| Q192T | hCoV-19/USA/OR-TRACE-BENT-011022-1536/2022 | EPI_ISL_12934316 | 2022-01-10 |
| Q192T | hCoV-19/USA/OR-TRACE-BENT-012421-725/2021 | EPI_ISL_6262416 | 2021-01-24 |
| Q192T | hCoV-19/USA/OR-TRACE-BENT-012421-737/2021 | EPI_ISL_6262422 | 2021-01-24 |
| Q192T | hCoV-19/USA/OR-TRACE-BENT-050921-707/2021 | EPI_ISL_2404763 | 2021-05-09 |
| Q192T | hCoV-19/USA/OR-TRACE-BENT-110421-1423/2021 | EPI_ISL_12934242 | 2021-11-04 |
| Q192T | hCoV-19/USA/OR-TRACE-BENT-122221-1330/2021 | EPI_ISL_12658860 | 2021-12-22 |
| Q192T | hCoV-19/USA/OR-TRACE-BENT-122421-1514/2021 | EPI_ISL_12934294 | 2021-12-24 |
| Q192T | hCoV-19/USA/OR-TRACE-BENT-122621-1469/2021 | EPI_ISL_12934171 | 2021-12-26 |
| Q192T | hCoV-19/USA/OR-TRACE-LINC-012121-719/2021 | EPI_ISL_6262411 | 2021-01-21 |
| Q192T | hCoV-19/USA/OR-TRACE-LINC-012521-739/2021 | EPI_ISL_6262424 | 2021-01-25 |
| Q192T | hCoV-19/USA/OR-TRACE-LINN-081421-992/2021 | EPI_ISL_7493998 | 2021-08-14 |
| Q192T | hCoV-19/USA/OR-TRACE-LINN-081621-926/2021 | EPI_ISL_7493988 | 2021-08-16 |
| Q192T | hCoV-19/USA/OR-TRACE-LINN-090721-1107/2021 | EPI_ISL_7551922 | 2021-09-07 |
| Q192T | hCoV-19/USA/OR-TRACE-LINN-111921-1329/2021 | EPI_ISL_12658834 | 2021-11-19 |
| Q192T | hCoV-19/USA/OR-TRACE-LINN-112821-1343/2021 | EPI_ISL_12658828 | 2021-11-28 |
| Q192T | hCoV-19/USA/OR-TRACE-LINN-121521-1339/2021 | EPI_ISL_12658858 | 2021-12-15 |
| Q192T | hCoV-19/USA/PA-0158/2021 | EPI_ISL_10657892 | 2021-05-17 |
| Q192T | hCoV-19/USA/TX-HMH-96473/2022 | EPI_ISL_12108959 | 2022-04-04 |
| Q192T | hCoV-19/USA/TX-HMH-99822/2022 | EPI_ISL_13367274 | 2022-06-04 |
| Q192T | hCoV-19/USA/TX-HMH-99855/2022 | EPI_ISL_13367265 | 2022-06-06 |
| Q192T | hCoV-19/USA/TX-HMH-M-105907/2022 | EPI_ISL_13993042 | 2022-07-10 |
| Q192T | hCoV-19/USA/TX-HMH-M-106911/2022 | EPI_ISL_14028264 | 2022-07-12 |
| Q192T | hCoV-19/USA/TX-HMH-MCoV-23104/2021 | EPI_ISL_1077634 | 2021-01-12 |
| Q192T | hCoV-19/USA/TX-HMH-MCoV-23787/2021 | EPI_ISL_1078317 | 2021-01-15 |

| | | | |
|---|---|---|---|
| Q192T | hCoV-19/USA/TX-HMH-MCoV-24629/2021 | EPI_ISL_1079211 | 2021-01-18 |
| Q192T | hCoV-19/USA/TX-HMH-MCoV-25306/2021 | EPI_ISL_1079935 | 2021-01-20 |
| Q192T | hCoV-19/USA/TX-HMH-MCoV-26172/2021 | EPI_ISL_1235952 | 2021-03-05 |
| Q192T | hCoV-19/USA/TX-HMH-MCoV-26864/2021 | EPI_ISL_1236645 | 2021-02-24 |
| Q192T | hCoV-19/USA/TX-HMH-MCoV-26887/2021 | EPI_ISL_1236668 | 2021-02-24 |
| Q192T | hCoV-19/USA/TX-HMH-MCoV-27095/2021 | EPI_ISL_1236876 | 2021-02-22 |
| Q192T | hCoV-19/USA/TX-HMH-MCoV-30408/2020 | EPI_ISL_1304961 | 2020-12-21 |
| Q192T | hCoV-19/USA/TX-HMH-MCoV-31346/2021 | EPI_ISL_2189950 | 2021-03-08 |
| Q192T | hCoV-19/USA/TX-HMH-MCoV-33939/2020 | EPI_ISL_2202168 | 2020-12-29 |
| Q192T | hCoV-19/USA/TX-HMH-MCoV-35235/2020 | EPI_ISL_2220396 | 2020-08-10 |
| Q192T | hCoV-19/USA/TX-HMH-MCoV-36070/2020 | EPI_ISL_2205920 | 2020-12-14 |
| Q192T | hCoV-19/USA/TX-HMH-MCoV-37634/2020 | EPI_ISL_2210662 | 2020-12-03 |
| Q192T | hCoV-19/USA/TX-HMH-MCoV-37969/2020 | EPI_ISL_2213364 | 2020-12-11 |
| Q192T | hCoV-19/USA/TX-HMH-MCoV-38455/2021 | EPI_ISL_2211824 | 2021-03-30 |
| Q192T | hCoV-19/USA/TX-HMH-MCoV-39560/2020 | EPI_ISL_2221626 | 2020-08-08 |
| Q192T | hCoV-19/USA/TX-HMH-MCoV-39599/2020 | EPI_ISL_2221740 | 2020-08-09 |
| Q192T | hCoV-19/USA/TX-HMH-MCoV-39903/2020 | EPI_ISL_2222655 | 2020-08-05 |
| Q192T | hCoV-19/USA/TX-HMH-MCoV-44165/2021 | EPI_ISL_2225403 | 2021-04-30 |
| Q192T | hCoV-19/USA/TX-HMH-MCoV-44932/2020 | EPI_ISL_5085326 | 2020-07-27 |
| Q192T | hCoV-19/USA/TX-HMH-MCoV-45207/2020 | EPI_ISL_5085630 | 2020-07-24 |
| Q192T | hCoV-19/USA/TX-HMH-MCoV-45319/2020 | EPI_ISL_5085794 | 2020-07-24 |
| Q192T | hCoV-19/USA/TX-HMH-MCoV-45428/2021 | EPI_ISL_5070007 | 2021-05-03 |
| Q192T | hCoV-19/USA/TX-HMH-MCoV-46381/2020 | EPI_ISL_5087116 | 2020-07-23 |
| Q192T | hCoV-19/USA/TX-HMH-MCoV-46745/2021 | EPI_ISL_5070771 | 2021-05-20 |
| Q192T | hCoV-19/USA/TX-HMH-MCoV-47305/2021 | EPI_ISL_5072669 | 2021-06-04 |
| Q192T | hCoV-19/USA/TX-HMH-MCoV-47879/2020 | EPI_ISL_5343126 | 2020-07-17 |
| Q192T | hCoV-19/USA/TX-HMH-MCoV-48347/2020 | EPI_ISL_5343441 | 2020-07-14 |
| Q192T | hCoV-19/USA/TX-HMH-MCoV-49818/2020 | EPI_ISL_6948520 | 2020-07-16 |
| Q192T | hCoV-19/USA/TX-HMH-MCoV-49984/2021 | EPI_ISL_6948691 | 2021-07-16 |
| Q192T | hCoV-19/USA/TX-HMH-MCoV-50579/2020 | EPI_ISL_6949388 | 2020-07-17 |
| Q192T | hCoV-19/USA/TX-HMH-MCoV-56909/2021 | EPI_ISL_5344478 | 2021-09-19 |
| Q192T | hCoV-19/USA/TX-HMH-MCoV-58774/2021 | EPI_ISL_5346405 | 2021-08-26 |
| Q192T | hCoV-19/USA/TX-HMH-MCoV-59355/2021 | EPI_ISL_5347013 | 2021-08-27 |
| Q192T | hCoV-19/USA/TX-HMH-MCoV-60114/2021 | EPI_ISL_5372207 | 2021-08-31 |
| Q192T | hCoV-19/USA/TX-HMH-MCoV-60498/2021 | EPI_ISL_5373359 | 2021-09-01 |

| | | | |
|---|---|---|---|
| Q192T | hCoV-19/USA/TX-HMH-MCoV-60851/2021 | EPI_ISL_5374503 | 2021-09-02 |
| Q192T | hCoV-19/USA/TX-HMH-MCoV-60900/2021 | EPI_ISL_5374614 | 2021-09-02 |
| Q192T | hCoV-19/USA/TX-HMH-MCoV-60985/2021 | EPI_ISL_5374817 | 2021-09-06 |
| Q192T | hCoV-19/USA/TX-HMH-MCoV-61235/2021 | EPI_ISL_5376075 | 2021-09-03 |
| Q192T | hCoV-19/USA/TX-HMH-MCoV-61342/2021 | EPI_ISL_5376878 | 2021-09-03 |
| Q192T | hCoV-19/USA/TX-HMH-MCoV-61369/2021 | EPI_ISL_5376944 | 2021-09-03 |
| Q192T | hCoV-19/USA/TX-HMH-MCoV-62002/2021 | EPI_ISL_5379263 | 2021-09-07 |
| Q192T | hCoV-19/USA/TX-HMH-MCoV-62411/2021 | EPI_ISL_5380759 | 2021-09-09 |
| Q192T | hCoV-19/USA/TX-HMH-MCoV-62858/2021 | EPI_ISL_5381611 | 2021-09-11 |
| Q192T | hCoV-19/USA/TX-HMH-MCoV-62910/2021 | EPI_ISL_5382067 | 2021-09-10 |
| Q192T | hCoV-19/USA/TX-HMH-MCoV-65018/2021 | EPI_ISL_5384785 | 2021-09-24 |
| Q192T | hCoV-19/USA/TX-HMH-MCoV-66071/2021 | EPI_ISL_6948931 | 2021-10-06 |
| Q192T | hCoV-19/USA/TX-HMH-MCoV-67395/2021 | EPI_ISL_7081417 | 2021-11-03 |
| Q192T | hCoV-19/USA/TX-HMH-MCoV-67683/2021 | EPI_ISL_7224338 | 2021-11-07 |
| Q192T | hCoV-19/USA/TX-HMH-MCoV-71170/2021 | EPI_ISL_9198712 | 2021-12-20 |
| Q192T | hCoV-19/USA/TX-HMH-MCoV-73230/2022 | EPI_ISL_9200168 | 2022-01-02 |
| Q192T | hCoV-19/USA/TX-HMH-MCoV-73737/2022 | EPI_ISL_9201012 | 2022-01-03 |
| Q192T | hCoV-19/USA/TX-HMH-MCoV-75569/2022 | EPI_ISL_9204861 | 2022-01-05 |
| Q192T | hCoV-19/USA/TX-HMH-MCoV-79765/2022 | EPI_ISL_10821855 | 2022-01-13 |
| Q192T | hCoV-19/USA/TX-HMH-MCoV-79871/2022 | EPI_ISL_10822946 | 2022-01-13 |
| Q192T | hCoV-19/USA/TX-HMH-MCoV-80701/2022 | EPI_ISL_10823749 | 2022-01-14 |
| Q192T | hCoV-19/USA/TX-HMH-MCoV-80766/2022 | EPI_ISL_10823660 | 2022-01-14 |
| Q192T | hCoV-19/USA/TX-HMH-MCoV-80807/2022 | EPI_ISL_10823392 | 2022-01-14 |
| Q192T | hCoV-19/USA/TX-HMH-MCoV-82352/2022 | EPI_ISL_10834833 | 2022-01-17 |
| Q192T | hCoV-19/USA/TX-HMH-MCoV-83045/2022 | EPI_ISL_10833303 | 2022-01-19 |
| Q192T | hCoV-19/USA/TX-HMH-MCoV-83861/2022 | EPI_ISL_10830733 | 2022-01-20 |
| Q192T | hCoV-19/USA/TX-HMH-MCoV-84333/2022 | EPI_ISL_10829376 | 2022-01-22 |
| Q192T | hCoV-19/USA/TX-HMH-MCoV-86535/2022 | EPI_ISL_10829142 | 2022-01-24 |
| Q192T | hCoV-19/USA/TX-HMH-MCoV-86658/2022 | EPI_ISL_10828542 | 2022-01-26 |
| Q192T | hCoV-19/USA/TX-HMH-MCoV-87806/2022 | EPI_ISL_10828848 | 2022-01-31 |
| Q192T | hCoV-19/USA/TX-HMH-MCoV-87848/2022 | EPI_ISL_10828853 | 2022-01-31 |
| Q192T | hCoV-19/USA/TX-HMH-MCoV-91190/2022 | EPI_ISL_10834039 | 2022-02-12 |
| Q192T | hCoV-19/USA/TX-HMH-MCoV-92016/2022 | EPI_ISL_10835226 | 2022-02-16 |
| Q192T | hCoV-19/USA/TX-HMH-MCoV-92028/2022 | EPI_ISL_10835234 | 2022-02-16 |
| Q192T | hCoV-19/USA/TX-HMH-MCoV-92076/2022 | EPI_ISL_10835085 | 2022-02-16 |

| | | | |
|---|---|---|---|
| Q192T | hCoV-19/USA/TX-HMH-MCoV-92115/2022 | EPI_ISL_10834666 | 2022-02-17 |
| Q192T | hCoV-19/USA/TX-HMH-MCoV-92116/2022 | EPI_ISL_10834665 | 2022-02-16 |
| Q192T | hCoV-19/USA/TX-HMH-MCoV-9302/2020 | EPI_ISL_1203772 | 2020-07-03 |
| Q192T | hCoV-19/USA/TX-HMH-MCoV-95505/2022 | EPI_ISL_11007858 | 2022-03-05 |
| Q192T | hCoV-19/USA/TX-HMH-MCoV-97139/2022 | EPI_ISL_12710048 | 2022-05-03 |
| Q192T | hCoV-19/USA/TX-HMH-MCoV-99078/2022 | EPI_ISL_13270273 | 2022-05-31 |
| Q192T | hCoV-19/USA/TX-HMH-MCoV-99449/2022 | EPI_ISL_13269991 | 2022-06-02 |

### References


(1) Zhao, Y., Fang, C., Zhang, Q., Zhang, R., Zhao, X., Duan, Y., Wang, H., Zhu, Y., Feng, L., Zhao, J., Shao, M., Yang, X., Zhang, L., Peng, C., Yang, K., Ma, D., Rao, Z., and Yang, H. (2021) Crystal structure of SARS-CoV-2 main protease in complex with protease inhibitor PF-07321332, *Protein Cell 13*, 689–693.

(2) Zhou, Y., Gammeltoft, K. A., Ryberg, L. A., Pham, L. V., Fahnøe, U., Binderup, A., Hernandez, C. R. D., Offersgaard, A., Fernandez-Antunez, C., Peters, G. H. J., Ramirez, S., Bukh, J., and Gottwein, J. M. (2022) Nirmatrelvir resistant SARS-CoV-2 variants with high fitness *in vitro*, *bioRxiv*, 2022.2006.2006.494921.

(3) Jochmans, D., Liu, C., Donckers, K., Stoycheva, A., Boland, S., Stevens, S. K., De Vita, C., Vanmechelen, B., Maes, P., Trüeb, B., Ebert, N., Thiel, V., De Jonghe, S., Vangeel, L., Bardiot, D., Jekle, A., Blatt, L. M., Beigelman, L., Symons, J. A., Raboisson, P., Chaltin, P., Marchand, A., Neyts, J., Deval, J., and Vandyck, K. (2022) The substitutions L50F, E166A and L167F in SARS-CoV-2 $3CL^{pro}$ are selected by a protease inhibitor *in vitro* and confer resistance to nirmatrelvir, *bioRxiv*, 2022.2006.2007.495116.

(4) Hu, Y., Lewandowski, E. M., Tan, H., Morgan, R. T., Zhang, X., Jacobs, L. M. C., Butler, S. G., Mongora, M. V., Choy, J., Chen, Y., and Wang, J. (2022) Naturally occurring mutations of SARS-CoV-2 main protease confer drug resistance to nirmatrelvir, *bioRxiv*, 2022.2006.2028.497978.

(5) Heilmann, E., Costacurta, F., Volland, A., and von Laer, D. (2022) SARS-CoV-2 $3CL^{pro}$ mutations confer resistance to Paxlovid (nirmatrelvir/ritonavir) in a VSV-based, non-gain-of-function system, *bioRxiv*, 2022.2007.2002.495455.

(6) de Oliveira, V. M., Ibrahim, M. F., Sun, X., Hilgenfeld, R., and Shen, J. (2022) H172Y mutation perturbs the $S_1$ pocket and nirmatrelvir binding of SARS-CoV-2 main protease through a nonnative hydrogen bond, *bioRxiv*, 2022.2007.2031.502215.

(7) Iketani, S., Mohri, H., Culbertson, B., Hong, S. J., Duan, Y., Luck, M. I., Annavajhala, M. K., Guo, Y., Sheng, Z., Uhlemann, A.-C., Goff, S. P., Sabo, Y., Yang, H., Chavez, A., and Ho, D. D. (2022) Multiple pathways for SARS-CoV-2 resistance to nirmatrelvir, *bioRxiv*, 2022.2008.2007.499047.

(8) Moghadasi, S. A., Heilmann, E., Moraes, S. N., Kearns, F. L., von Laer, D., Amaro, R. E., and Harris, R. S. (2022) Transmissible SARS-CoV-2 variants with resistance to clinical protease inhibitors, *bioRxiv*, 2022.2008.2007.503099.

(9) Zhou, P., Yang, X.-L., Wang, X.-G., Hu, B., Zhang, L., Zhang, W., Si, H.-R., Zhu, Y., Li, B., Huang, C.-L., Chen, H.-D., Chen, J., Luo, Y., Guo, H., Jiang, R.-D., Liu, M.-Q., Chen, Y., Shen, X.-R., Wang, X., Zheng, X.-S., Zhao, K., Chen, Q.-J., Deng, F., Liu, L.-L., Yan, B., Zhan, F.-X., Wang, Y.-Y., Xiao, G.-F., and Shi, Z.-L. (2020) A pneumonia outbreak associated with a new coronavirus of probable bat origin, *Nature 579*, 270–273.

(10) Khare, S., Gurry, C., Freitas, L., B Schultz, M., Bach, G., Diallo, A., Akite, N., Ho, J., Tc Lee, R., Yeo, W., Core Curation Team, G., and Maurer-Stroh, S. (2021) GISAID's role in pandemic response, *China CDC Weekly 3*, 1049–1051.
